# Remote homology and functional genetics unmask deeply preserved Scm3/HJURP orthologs in metazoans

**DOI:** 10.64898/2026.03.04.709615

**Authors:** Jeremy A. Hollis, Jason A. Stonick, Irini Topalidou, Janet M. Young, Cecilia B. Moens, Nicolas J. Lehrbach, Melody G. Campbell, Harmit S. Malik

**Affiliations:** Molecular and Cellular Biology Graduate Program, University of Washington, Seattle, WA 98195 USA; Division of Basic Sciences; Howard Hughes Medical Institute, Fred Hutchinson Cancer Center; Seattle, WA 98109, USA

**Keywords:** *Drosophila melanogaster*, *Caenorhabditis elegans*, *Danio rerio*, coelacanth, horseshoe crab, Hymenoptera, AlphaFold, CenH3, CENPA, HJURP, centromere, remote homology

## Abstract

In most animals and fungi, centromere identity and function depend on the Scm3/HJURP chaperone, which deposits CENPA at centromeres. However, Scm3/HJURP orthologs appeared to be missing in insects, nematodes, many vertebrates, and other metazoans, suggesting radical chaperone replacement in these lineages. Here, we combine remote homology detection, AlphaFold-based structural modeling, and functional genetics in zebrafish and *Caenorhabditis elegans* to identify previously unknown Scm3/HJURP orthologs that localize to centromeres and whose loss causes catastrophic mitotic failure. We further show that Drosophila CAL1, long considered a functional analog, is instead a highly diverged Scm3/HJURP ortholog. Despite rapid primary-sequence divergence, predicted and known structures reveal a broadly conserved CENPA-H4-binding scm3 fold across fungi, vertebrates, nematodes, insects, and most metazoans. Our work demonstrates how rapid divergence can obscure the broad conservation of essential centromere machinery and provides a generalizable strategy for unmasking missing orthologs.

**Teaser:** Animals encode a rapidly evolving, essential cell cycle gene previously thought to be absent.

## Introduction

In most eukaryotes, accurate chromosome inheritance during cell division depends on the precise assembly of centromeric chromatin, a process orchestrated by the deposition of the centromere-specific histone H3 (CenH3) variant, CENPA(*1*). In fungi and mammals, this deposition is facilitated by a specialized histone chaperone(*2*) (Fig. S1A). *S. cerevisiae* Scm3(*3–5*) and human HJURP(*6–8*) were independently identified for their roles in CENPA deposition through genetic and biochemical studies. Subsequent evolutionary analyses based on remote homology detection identified fungal Scm3 and animal HJURP proteins as distantly related orthologs(*9*). Intriguingly, this orthology could not be directly established using traditional methods such as reciprocal BLAST genome queries. Instead, the identification of a choanoflagellate Scm3/HJURP protein, with homology to both fungal and animal proteins, provided a critical evolutionary intermediate that enabled the successful identification of orthology between the fungal Scm3 and animal HJURP proteins. Subsequently, several structural and functional studies demonstrated that Scm3(*10*, *11*) and HJURP(*12*) share a common ‘scm3 domain’(*13*) (we refer to the domain as <u>s</u>cm3 to distinguish it from the <u>S</u>cm3 protein in yeast and choanoflagellates) that mediates the specific recognition and assembly of CENPA-containing nucleosomes at centromeres, highlighting orthology of these proteins despite divergence of their primary sequences. This functional conservation underscores the significance of Scm3/HJURP-mediated CENPA loading and stabilization(*14*, *15*) in eukaryotic chromosome segregation, dating back to at least the common ancestor of fungi and animals. In contrast, plants appear to utilize a distinct chaperone, nuclear autoantigenic sperm protein (NASP), to deposit CENH3 in *Arabidopsis thaliana* and other land plants(*16*, *17*).

At the molecular level, the Scm3 and HJURP chaperones recognize the CENPA-Targeting Domain (CATD)(*18*), which encompasses the L1 loop and the α2 helix of the CENPA histone fold domain (Fig. S1B). The unique molecular features of the CENPA CATD are both necessary and sufficient for Scm3/HJURP to distinguish it from other H3 variants and to deposit CENPA strictly at the centromere, thereby making the scm3 domain critical for chaperone specificity(*19*). The scm3 domain is comprised of a structurally conserved ∼25aa α-helix followed by a ∼10aa β-strand that runs antiparallel to the CENPA α2 helix and L1 loop, respectively(*10*, *12*). Most of HJURP lacks experimental structural information and is predicted to be largely disordered. Scm3/HJURP proteins carrying CENPA/H4 dimers are recruited to the centromere via two parallel pathways: the first utilizes the polo-like kinase-licensed Mis18 complex to recruit HJURP(*20–22*), while the second involves HJURP binding directly to the inner kinetochore protein CENPC(*23*, *24*). These pathways appear to be broadly functionally conserved across the metazoan tree(*25–28*).

Despite its ancient origins, the evolutionary history of the Scm3/HJURP chaperone remains unclear in many animal lineages. Although sequence profile analyses and sensitive homology searches have identified distant relatives of Scm3 and HJURP in some lineages(*9*), orthologs appear to be missing in several major animal groups, especially in fish, insects, and nematodes(*29*) (Fig. S2). This is particularly puzzling, as most of these groups still possess canonical CENPA-dependent centromeres(*30*), indicating that the need for a CENPA chaperone persists, even if the specific chaperone might differ. The absence of detectable Scm3/HJURP homologs in these species has been taken as evidence for the independent evolution of distinct centromeric chaperones or of entirely different centromere assembly processes(*9*, *31*).

The case of insects, especially *Drosophila*, has been central to this puzzle. Genetic and cell biological studies have unambiguously identified the CAL1 protein as the CENPA chaperone in *Drosophila melanogaster*(*32–34*) and other dipteran species(*35*), where it is thought to play a role similar to that of HJURP/Scm3 in CENPA deposition. However, CAL1 shows no obvious sequence similarity to Scm3/HJURP, leading to its classification as a functional analog rather than a true ortholog(*31*). This distinction is based on the failure to detect homology to the conserved scm3 domain using standard sequence comparison tools. Nonetheless, structural studies have shown that CAL1 does contain an ‘scm3 domain’-like motif, which interacts with CENPA molecules similarly to Scm3/HJURP proteins(*36*). Therefore, based on current evidence, it remains unclear whether CAL1 is a true ortholog whose orthology is obscured by rapid divergence or an example of convergent evolution, in which a new protein has independently gained the ability to chaperone CENPA.

Unlike insects, nematodes such as *Caenorhabditis elegans* appear to lack a recognizable CENPA chaperone that is either orthologous to Scm3/HJURP or a clear functional equivalent, such as CAL1(*25*, *37*). Instead, recent research suggests that the exceptionally long, helical CENPA N-terminal tail, combined with the centromere protein KNL-2, may have usurped chaperone functions in centromeric chromatin assembly, potentially eliminating the need for a dedicated CENPA chaperone(*25*, *27*). In support of this model, genetic and cytological studies have shown that KNL-2, which lacks homology to Scm3/HJURP, is essential for CENPA loading and kinetochore formation in nematodes(*25*, *27*). This, combined with their holocentric architecture, suggests that *C. elegans* and possibly other nematodes may have evolved a fundamentally different mechanism for centromere specification and maintenance, highlighting the remarkable possibility that the machinery involved in depositing centromeric histones could be highly adaptable across evolution.

Here, we revisit this important question using both classical genetic approaches and recent advances in remote homology detection(*38*, *39*), which incorporate sequence comparisons and structural predictions, along with recently sequenced genomes (Fig. S3). From these analyses, we demonstrate that most metazoan clades have preserved an orthologous centromeric chaperone characterized by the conserved scm3 domain. For example, we identified previously undetected scm3 domain-containing proteins in deuterostomes, including teleost fish, nematodes, and a wide range of arthropods. Using genetic perturbations and cytological analyses, we functionally validate our identification of Scm3/HJURP orthologs in zebrafish (*Danio rerio*) and nematodes (*C. elegans*), demonstrating that these putative orthologs localize to centromeres and that their loss leads to catastrophic mitotic defects. We also trace the arthropod ancestry of the scm3 domain, revealing that *Drosophila* CAL1 is a genuine Scm3/HJURP ortholog rather than a functional analog. Structural modeling using AlphaFold(*40*, *41*) predicts that even highly diverged Scm3 domains from species such as zebrafish and nematodes adopt a conserved fold and interact with CENPA/H4 in a manner similar to that of their human and fungal counterparts. Our discovery of scm3 domain-containing proteins in basal arthropods, non-dipteran insects, nematodes, and other early-branching metazoans suggests that the ancestral metazoan and most metazoan lineages possess CENPA chaperones, which include an scm3 domain. Our findings comprehensively revise our understanding of centromeric histone chaperone evolution in animals, highlight the limitations of standard homology detection methods in the face of extreme sequence divergence, and offer general approaches to overcome these limitations.

## Results

### HJURP orthologs are broadly conserved across deuterostomes, including teleosts

Since the initial study identifying orthology between human HJURP and yeast Scm3(*9*), several additional high-quality genome assemblies across the metazoan tree have become available(*42*). These genomes have provided more power to identify distant homology between clades. We employed an iterative BLAST (PSI-BLAST)(*43*) approach (Fig. S3), which leverages repeated searches with new hits incorporated at each iteration, to identify remote homologs of full-length human HJURP in teleosts (ray-finned fish). Consistent with previous efforts, we were initially unsuccessful. However, we could confidently identify an HJURP-like protein in the genome of *Latimeria chalumnae* (West Indian Ocean coelacanth)(*44*), a lobe-finned fish, despite this sequence lacking the canonical scm3 domain. Instead, the coelacanth protein was identified by homology with human HJURP residues outside the scm3 domain. Despite lacking an scm3 domain, subsequent iterative searches using the putative coelacanth HJURP sequence successfully identified previously uncharacterized proteins across deuterostomes, including echinoderms, teleost fish, and cephalochordates (Fig. 1A). In most deuterostome genomes, only a single strong hit was identified. We generated a MUSCLE(*45*) alignment of these putative HJURP homologs from representative tetrapods, teleosts, and echinoderms, revealing a conserved scm3 domain across all identified sequences. Our results are supported by the InterPro database(*46*), which also lists several of these putative deuterostome HJURP proteins as containing a *bona fide* scm3 domain. Phylogenetic analyses of these scm3 domains were generally congruent with the deuterostome species tree(*47*) (Fig. 1B, Fig. S4A), consistent with the hypothesis that a monophyletic, broadly retained common HJURP ancestor is present among all deuterostomes.

**Figure 1.**
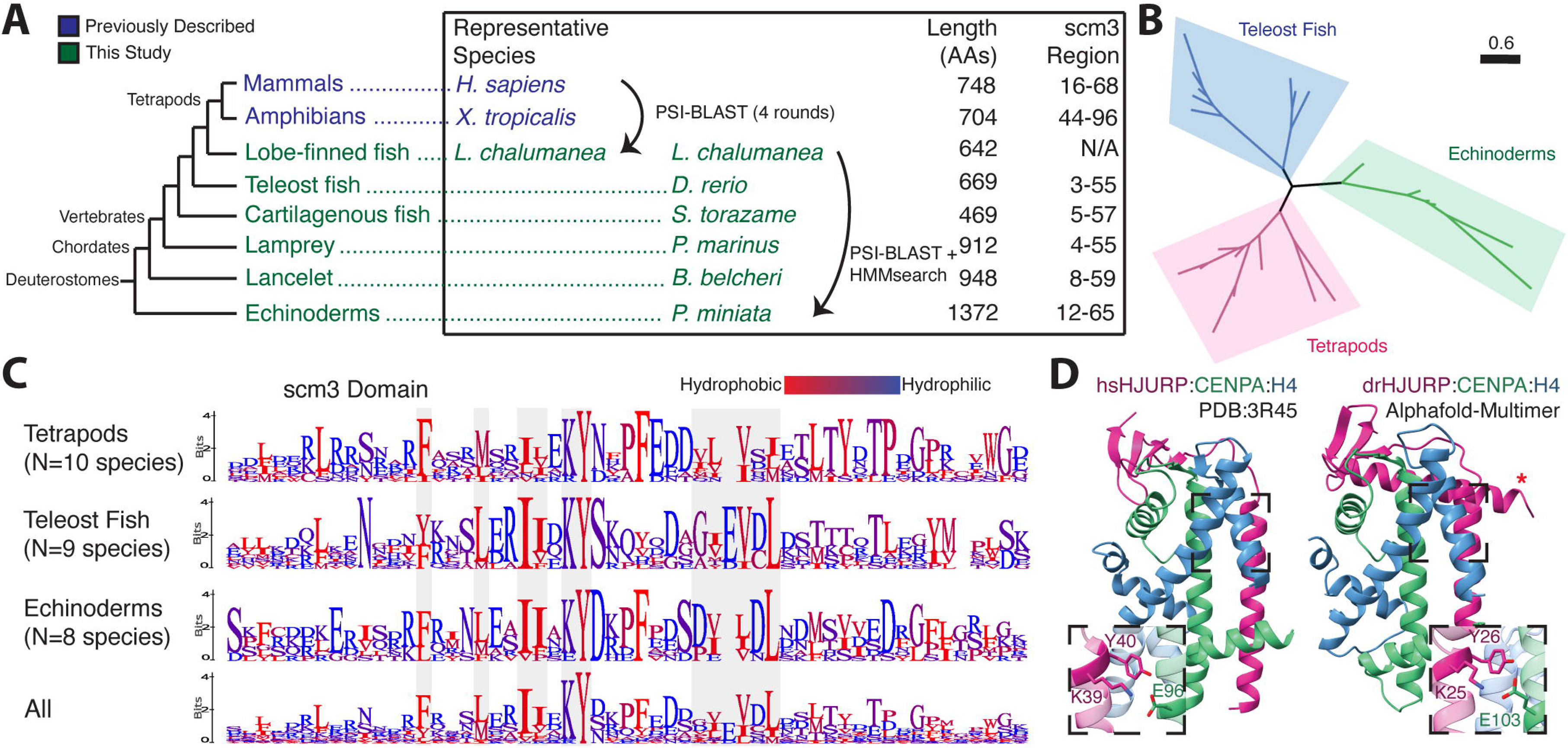
The centromeric chaperone HJURP predates tetrapods. **A.** A phylogeny (not to scale) of deuterostome putative HJURP proteins. Deuterostome HJURPs have high variation in length and composition, sharing only a short N-terminal scm3 domain. **B.** Phylogenetic analysis shows that putative deuterostome scm3 domains roughly follow the species tree, consistent with a monophyletic origin of HJURP. **C.** Logo plots of the scm3 domain for representative clades within deuterostomes. scm3 domains were extracted, aligned together, and then separated into distinct clades for the purposes of visualization. **D.** The *Homo sapiens* crystal structure (PDB:3R45, left) and AlphaFold-Multimer prediction of the *Danio rerio* HJURP-CENPA-H4 complex (right) show general structural similarities. The KY motif (insets) interacts with a conserved glutamate on the CENPA 122 helix.

Overall, these sequences showed little homology among deuterostomes, particularly outside the scm3 domain (9.91% average whole-protein identity), making whole-protein alignments intractable and uninformative (Fig. S4B). Even within the scm3 domain, there is relatively little sequence conservation (27.26% shared identity) compared to what would be expected of an essential domain in an essential protein(*48*). Logo plots of scm3 domains from representative clades show only a few broadly conserved amino acids (Fig. 1C). Among these is the previously identified KY dipeptide, which is almost universally conserved among deuterostomes, with only select conservative substitutions at the lysine residue. Hydrophobic amino acids spaced approximately 2-4 positions apart also appear to be important for the scm3 domain (Fig. 1C); this regular spacing is consistent with the alpha-helical structure of the scm3 domain that runs antiparallel to the l12 helix in the CENPA histone fold domain (Fig. S1B). An HHpred(*49*) E-value of 2.30e-04 supports this finding of structural homology (Table S1). HHpred values were also used as an additional validation metric for subsequent searching.

Conversely, while these general patterns define the scm3 domain, it appears to exhibit remarkable sequence plasticity even within deuterostomes. For example, a threonine within the “TLTY” motif was previously thought to be widely conserved across opisthokonts (encompassing fungi, choanoflagellates, and metazoa)(*9*, *13*), but shows only low conservation among these representative deuterostome clades. There is marked diversity in hydrophobic residues helical within and among clades (Fig. 1C). Furthermore, some residues are conserved in a lineage-specific manner. For example, the consensus residue immediately following the KY is conserved within, but differs across, representative lineages. Such variation may reflect lineage-specific coevolution between HJURP and the CENPA/H4 histone dimers. Although the scm3 domain appears to be the single shared molecular feature across HJURP proteins in deuterostomes, tetrapods and teleost fish share an additional motif immediately upstream of the DNA-binding HMD region (Fig. S1B) as well as moderate homology within the previously identified(*22*) Mis18-binding R2 region (Fig. S4C). This region of homology is not shared with echinoderms, suggesting that it arose and became entrenched for functional purposes in the common ancestor of tetrapods and teleosts.

Identification of deuterostome HJURPs confirmed that the scm3 domain is the definitive conserved feature of HJURP proteins in deuterostomes. Therefore, we hypothesized that the absence of the scm3 domain in the original coelacanth *L. chalumnae* HJURP protein must have arisen from an assembly error or gap in the coelacanth genome at this site. To further test this hypothesis, we used Logan Search(*50*) with the *L. chalumnae* gene sequence to query the Sequence Read Archive for homology to this gene. We found a close match to a transcript from the testes of the closely related *L. menadoensis* (Indonesian coelacanth)(*51*) that, when translated, extended the N-terminus of this protein to include an intact scm3 domain (Fig. S4D). Consistent with our assembly junction hypothesis, we could align two different *L. chalumnae* contigs to the bases 1-92 and 157-2691, respectively, of the *L. menadoensis* transcript. However, a contig gap spanning bases 93-156 rendered the sequence of the *L. chalumnae* HJURP homolog and its scm3 domain incomplete (Fig. S4D). Overall, our results indicated the scm3 domain is invariant in HJURP orthologs and provides the most conclusive signal for HJURP ortholog identification.

Our remote homology analyses revealed that a previously uncharacterized gene from zebrafish (*Danio rerio*) (ZFIN Gene ID: ZDB-GENE-030804-4; name: *si:dkeyp-117h8.*4(*52*)) encodes an HJURP ortholog, which we refer to as drHJURP. To gain further insight into the structural constraints that might shape the scm3 domain’s evolution in deuterostomes, we performed AlphaFold-multimer analyses of drHJURP in complex with the zebrafish CENPA-H4 dimer. Despite the low sequence homology to scm3 from human HJURP, AlphaFold-multimer predicted with high confidence a protein complex very similar in structure to the *Homo sapiens* crystal structure (Fig. 1D). Features broadly conserved in vertebrate scm3 domains include the KY motif (Fig. 1D) and a downstream motif comprised of acidic (E/D) and short hydrophobic residues (I/L/V) in a repeating pattern which forms the secondary beta sheet structure. These residues appear to perform orthologous functions in the case of human and zebrafish complexes. However, six CENPA residues essential for specificity in the human HJURP-CENPA interaction(*53*) are entirely different in *Danio rerio*, as are the complementary HJURP residues (Fig. S4E). Nevertheless, the essential pocket cradling CENPA’s specificity-defining CENPA-Targeting Domain (CATD)(*18*) is considerably similar in shape, with a backbone RMSD of 0.71Å^2^ between the helix-sheet region of the human and predicted zebrafish structures, despite dramatic sequence differences. This AlphaFold prediction reveals the structural basis for sequence plasticity within the scm3 domain: the strongest constraints arise from entrenched hydrophobic interactions (*12*, *53*), which are maintained in the scm3 domain to complement the hydrophobic CENPA residues required for tight nucleosome formation at the centromere. Surprisingly, the AlphaFold prediction also included an additional helix C-terminal to the scm3 domain that also appears to contact the histone dimer (Fig. 1D). This feature is reminiscent of the *Drosophila* CAL1-CID-H4(*36*) crystal structure, which shows that further tertiary structure beyond the central scm3-CENPA parallel helices helps augment CAL1-CENPA interaction specificity.

### The zebrafish gene *si:dkeyp-117h8.4* encodes HJURP

To assess the centromeric function of drHJURP *in vivo*, we tested three critical attributes. We first asked whether drHJURP preferentially interacts with drCENPA rather than with canonical H3. To detect a direct interaction between drCENPA and drHJURP, we performed *in vitro* pull-down assays using recombinant proteins. These experiments revealed the hallmark preference of HJURP for CENPA/H4 over canonical H3/H4(*54*) (Fig. 2A). However, we note that this pulldown with drHJURP was qualitatively less efficient than those performed with orthologous tetrapod proteins, including human HJURP(*53–55*).

**Figure 2.**
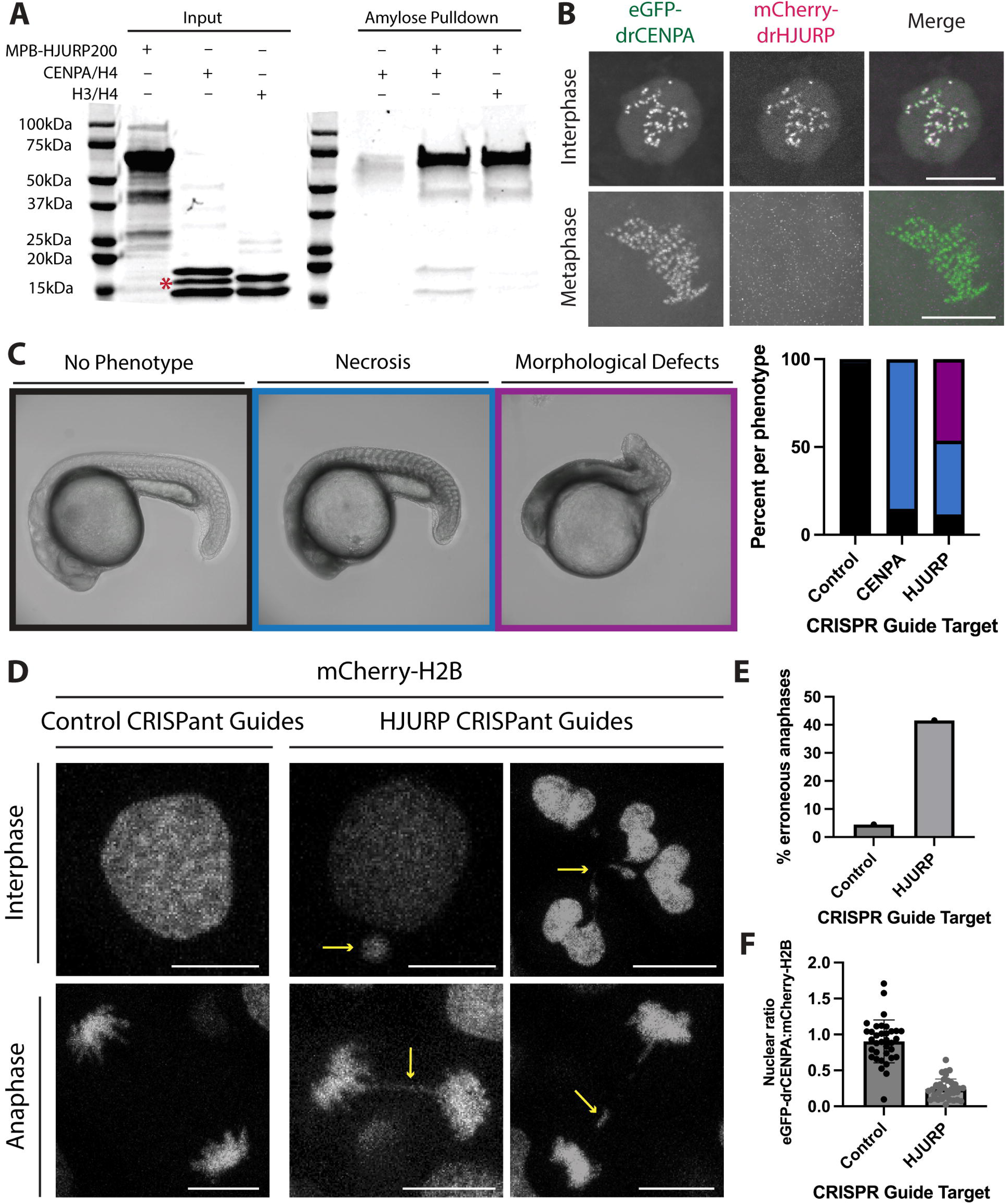
*Danio rerio* encodes an HJURP ortholog. A. Amylose pulldown assay with purified MBP-drHJURP and drCENPA/H4 or canonical H3/H4 tetramers. MBP-drHJURP interacts directly with drCENPA/H4 tetramers as suggested by co-precipitation of species at drCENPA and H4 sizes. Red asterisk indicates a non-histone contaminant. B. Still frames from timelapse series of cell divisions in *D. rerio* embryos expressing GFP-CENPA and mCherry-HJURP show cell cycle-specific colocalization of drHJURP and drCENPA. C. Morphological analysis of CRISPant zebrafish larvae with control (left representative image, N=96), CENPA (N=81), or HJURP (center and right representative images, N=138) guides. Severe morphological defects suggest HJURP is an essential gene. D. While control guide-injected embryonic cells generally segregate their chromosomes cleanly, HJURP CRISPant embryos have severe chromosome segregation defects, including frequently aberrant anaphases and fragmented nuclei. Scale bars are 5μm. E. Quantification of anaphase defects in control or HJURP CRISPant embryos. Numbers of anaphases are 49 for control guides, 105 for HJURP guides. F. Relative centromeric CENPA fluorescence in control CRISPant embryos (N=34 nuclei) compared to HJURP CRISPant embryos (N=36 nuclei) suggests a loss of centromeric CENPA in the absence of HJURP.

A second expectation is that drHJURP should localize to centromeres. To test this expectation, we co-injected zebrafish zygotes with *in vitro*-transcribed mRNAs encoding eGFP-tagged drCENPA (to mark centromeres) and mCherry-tagged drHJURP, then examined the localization of these tagged proteins via live imaging. We observed strong colocalization of drCENPA and drHJURP in embryonic nuclei (Fig. 2B). This colocalization was cell-cycle dependent: drHJURP was present at centromeres only in interphase nuclei (16/25 nuclei) and was completely absent in other phases of the cell cycle (Fig. 2B, Movie S1). In contrast, drCENPA localized to centromeres throughout the cell cycle. This localization pattern mirrors that of human HJURP, which replenishes CENPA at the telophase-to-G1 transition(*6*, *7*, *56*). The cell-cycle-dependent localization of drHJURP is also consistent with its expression being most strongly correlated with other cell-cycle components (Fig. S5A) in single-cell transcriptomics data(*57*, *58*) and *in situ* hybridization(*59*).

Finally, we tested the third expectation for drHJURP; given its fundamental role in CENPA deposition, we expected that drHJURP would be essential for early embryonic development(*60*). To test this possibility, we injected zebrafish embryos with Cas9 protein and CRISPR guide RNAs targeting drHJURP, drCENPA, or a scrambled negative control (Fig. S5B). We then examined CRISPant F0 embryos looking for indications of developmental impairment. While the control group was largely unimpaired, drHJURP CRISPant embryos displayed gross morphological defects and extensive signs of necrotic tissue at 24hpf (Fig. 2C). By comparison, drCENPA CRISPants displayed a comparatively milder, yet substantial phenotype. We hypothesize that the differences in phenotypic severity between the drHJURP and drCENPA CRISPants can be attributed to the exceptionally long half-life of the CENPA protein(*61*), together with its considerable maternal deposition. If the primary cause of necrosis in drHJURP and drCENPA CRISPant embryos is centromeric dysregulation, we hypothesized that we would observe signs of catastrophic cell division. Consistent with this expectation, we found extensive evidence of mitotic failure in drHJURP CRISPant embryos, including lagging and bridging chromosomes in anaphase cells as well as fractured nuclei and micronuclei in interphase cells (Fig. 2D-E). Furthermore, we find a ∼65% reduction of GFP-drCENPA intensity in drHJURP CRISPant embryos compared to scramble controls (Fig. 2F). Thus, our *in vivo* data indicate that the *D. rerio* gene *si:dkeyp-117h8.4* indeed encodes the long-missing zebrafish HJURP, validating the conclusions from our remote homology search.

### *Drosophila* CAL1 is a true HJURP ortholog

Based on current evidence, it is unclear whether the CAL1 CENPA chaperone identified in *Drosophila* and other *Diptera* represents a divergent Scm3/HJURP ortholog or whether it independently evolved to converge on HJURP/Scm3-like structure and function(*29*, *31*). The success of our remote homology detection strategy in identifying HJURP orthologs in deuterostome lineages led us to investigate whether we could distinguish between these two possibilities by extending our search to arthropods. To this end, we sought to identify broadly distributed arthropod proteins that contain scm3 domains. We generated different Hidden Markov Models (HMMs) via HMMer(*62–64*) either from deuterostome HJURP scm3 domains, yeast/choanoflagellate scm3 domains, or Dipteran CAL1 proteins, encompassing either just the scm3-like domain or the first 160 residues (Data S1). We used these different HMM models to search available arthropod genomes for putative Scm3/HJURP homologs.

To our surprise, we found a non-overlapping set of putative HJURP homologs using these strategies. For example, using the deuterostome or yeast/choanoflagellate HMMs, we were able to successfully identify scm3 domain-encoding genes from basal panarthropods, such as tardigrades and horseshoe crabs, suggesting that the scm3 domain was present in the arthropod common ancestor (Fig. 3A). However, this search did not initially reveal any homologs in insects. Conversely, using Dipteran-based HMMs, we identified proteins with scm3-like domains in *Diptera* and other insect clades, including *Hymenoptera* (bees, ants, wasps), *Coleoptera* (beetles), and *Orthoptera* (crickets), but not in other arthropods. However, by searching arthropod genomes using iteratively built HMMs combined with AlphaFold modeling, we identified representative scm3 domain-containing proteins in all major clades of arthropods (Fig. 3A). Scm3/HJURP orthologs from *Hymenoptera* were extremely useful evolutionary intermediates because they were found to be homologous to both deuterostome HJURP and insect CAL1 proteins, thus confirming the orthology between dipteran CAL1 and non-insect arthropod Scm3/HJURP proteins. These findings provide the most definitive evidence that *Drosophila* CAL1 is a true, albeit more divergent, ortholog of Scm3/HJURP.

**Figure 3.**
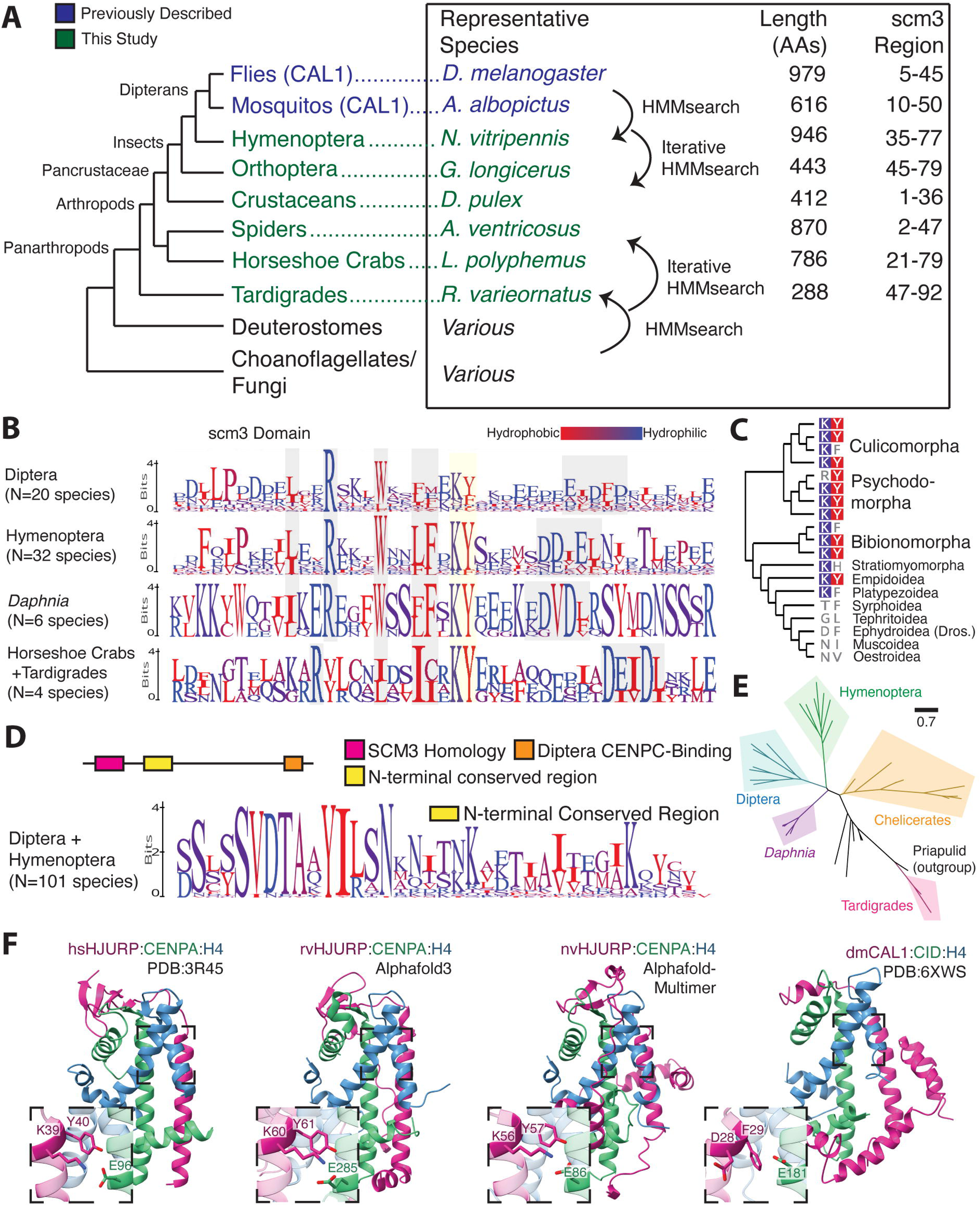
CAL1 is a true Scm3/HJURP ortholog. **A.** A schematic of the approach used to establish homology of the scm3 domain between basal arthropods and Dipteran CAL1. We find homologous scm3-like sequences across the arthropod lineage. As with deuterostomes, arthropod sequences are highly variable and share only the scm3 domain. **B.** Logo plots of the scm3 domain for clades within panarthropods suggest broadly conserved residue-level sequence features (gray and yellow highlighting) are present across arthropod scm3 orthologs. **C.** A Dipteran species phylogeny showing amino acids aligned to the conserved KY positions, suggests that the constraints on this motif were lost relatively recently in fly evolution. **D.** Sequence homology is shared beyond the scm3 domain between Hymenoptera and Diptera. A motif flanking the C-terminal side of the scm3 domain shares more homology between the orders than the scm3 domain itself does. **E.** A phylogenetic tree constructed using full length HJURP sequences from representative arthropod species generally follows the species tree, supporting a monophyletic origin of HJURP in arthropods. **F.** Structural predictions of representative tardigrade (rvHJURP) and Hymenopteran (nvHJURP) scm3 domains complexed with their respective species’ CENPA and H4 suggest a stepwise progression from the human HJURP structural fold to the *D. melanogaster* CAL1 fold.

Based on our more comprehensive repertoire of Scm3/HJURP orthologs in arthropods, we sought to understand why earlier efforts might have failed to establish CAL1 as a true HJURP ortholog. We focused on sequence features that make CAL1 appear distant from other proteins with scm3 domains. To our surprise, we found that, unlike deuterostomes, the scm3 domain is not the most conserved region of the CAL1 chaperone in *Diptera* (Fig. S6A). Rather, the scm3 domain shows comparably low sequence homology within dipterans compared to other arthropod lineages (Fig. 3B). A subset of Dipterans has also lost selective constraint on the definitive KY motif (Fig. 3C), although the orthologous DF residues in *Drosophila melanogaster* remain functionally important(*36*). This loss may be dipteran-specific, as the KY is still largely present in the hymenopteran insect clade (Fig. 3B), as is the acidic/short hydrophobic patch. Other features appear to be universal among insects, such as largely invariant tryptophan and arginine residues, which are also widely conserved in the *Daphnia* crustacean lineage. The regular spacing of conserved residues is also consistent with both deuterostome scm3 sequences and the expected parallel helical structure of the chaperone-histone complex(*36*). Despite these features, insect scm3 domain sequences are considerably diverged from those of early-branching arthropods (Fig. 3B). The relaxed sequence constraints on dipteran scm3 domains may explain why initial attempts failed to establish CAL1’s orthology with human HJURP, necessitating the requirement for evolutionary intermediates from other insects and other arthropods described here.

As support for the hypothesis of altered constraints on the scm3 domain in Diptera, we found that the region immediately C-terminal to the scm3 domain in *Diptera* and *Hymenoptera* shows generally higher sequence homology than the *Diptera* scm3 domain itself (Fig. S6A, Fig. 3D). This region may form additional contacts beyond the scm3 domain, although it lies just beyond the CAL1 fragment used for the crystal structure(*36*). These data together suggest the *Hymenoptera* CAL1 is a representative evolutionary intermediate between the ancestral HJURP and the dipteran CAL1 of monophyletic origin; phylogenetic analysis generally supports this placement (Fig. 3E, Fig. S7).

We employed AlphaFold-multimer(*41*) to predict conserved features of the putative *Nasonia vitripennis* (*Hymenoptera*) HJURP. We found that the predicted nvHJURP-CENPA-H4 structural topology is highly similar to the *Drosophila melanogaster* CAL1-CENPA-H4 crystal structure (Fig. 3F). Secondary and tertiary features of the complex appear to be conserved between insect clades. However, the nvHJURP prediction contains antiparallel beta strands with the CATD region of nvCENPA, which more closely resemble those in the *Homo sapiens* HJURP structure(*12*) but are absent from the *Drosophila* CAL1 crystal structure. Thus, structural predictions of *Nasonia* HJURP show intermediate features between basal arthropod HJURP and dipteran CAL1 sequences. This supports our hypothesis that constraints on *Drosophila* CAL1 are relaxed or altered relative to other Scm3/HJURP orthologs. These observations suggest a gradual evolutionary trajectory in which the enhanced function of domains C-terminal to the scm3 region in *Diptera* may have relaxed constraints on the scm3 domain in early dipterans. Such a change in constraint would have challenged prior efforts to establish orthology based solely on the scm3 domain. Further, if the hymenopteran prediction represents a structural intermediate, we would expect a more basal arthropod structure to more closely resemble the canonical human and yeast-like fold. Consistent with this expectation, the predicted HJURP-CENPA-H4 structure from the tardigrade *R. varieornatus* (sister clade to all arthropods) also shows features more consistent with the human protein crystal structure, such as a tighter antiparallel helix interface and an extended beta sheet region (Fig. 3F).

In addition to an altered scm3 domain and an additional domain immediately C-terminal to scm3 (Fig. 3B-D), *Drosophila* CAL1 proteins have additional unique evolutionary constraints not seen in other CENPA chaperones(*29*, *65*), including a C-terminal CENPC-binding domain whose function is thought to replace Mis18 binding present in tetrapod chaperones(*36*, *66*, *67*). In contrast to *Diptera*, CAL1/HJURPs in Hymenoptera show comparatively low conservation of their C-terminal regions, and there is little likeness between *Diptera* and *Hymenoptera* in this region (Fig. S6B). These findings suggest that the CAL1 CENPC-binding domain was acquired after the divergence of *Diptera* and *Hymenoptera*, further highlighting the plasticity of the centromeric machinery across the metazoan tree. Despite these idiosyncrasies, our findings trace the evolutionary trajectory of HJURP within arthropods and strongly support the hypothesis that CAL1 and HJURP proteins are genuine orthologs.

### Nematodes encode HJURP proteins

Another metazoan lineage that appears to lack Scm3/HJURP orthologs entirely is the nematode lineage. The holocentric nematode *C. elegans* has proven instrumental in advancing our understanding of the fundamental principles that govern the cell cycle, with detailed genetic and cell biological analyses identifying most of the currently known essential kinetochore components in nematodes(*68*). Nevertheless, neither previous genetic screens nor prior *in silico* and *in vitro* studies have successfully identified an HJURP ortholog in nematodes(*9*, *69*). Instead, the current model suggests that the exceptionally long N-terminal tail of the *C. elegans* CENPA homolog HCP-3 (hereafter called CENPA^HCP-3^) interacts uniquely with the centromeric chromatin assembly factor KNL-2, partially replacing HJURP function(*25*, *27*). However, this interaction alone is insufficient to explain how CENPA^HCP-3^ is stably recruited from the cytosol to the centromere.

Given this functional gap, we sought to determine if our remote homology approach could identify a *bona fide* HJURP in nematodes, including *C. elegans*. Using the scm3/HJURP HMM approach, which was successful in deuterostomes and arthropods, we were still unable to identify an HJURP ortholog in *C. elegans*. This lack of success appeared to confirm current models that nematodes might lack HJURP orthologs. However, our search criteria identified a region of minimal scm3 homology in the protein CAI4225225 from the nematode species *Auanema sp. JU1783*(*70*), which is only distantly related to *Caenorhabditis*. Using this sequence as a query, we performed iterative BLAST searches of nematode genomes, eventually revealing remote homology to an scm3 domain in *C. elegans* gene *nefr-1*(*71*) (Fig. 4A). As with deuterostomes, we found the phylogenetic placement of putative HJURP orthologs largely follows the nematode species tree(*72*), suggesting that the common ancestor of nematodes encoded a single HJURP ortholog that has been stably inherited across nematodes (Fig. 4B, S8A).

**Figure 4.**
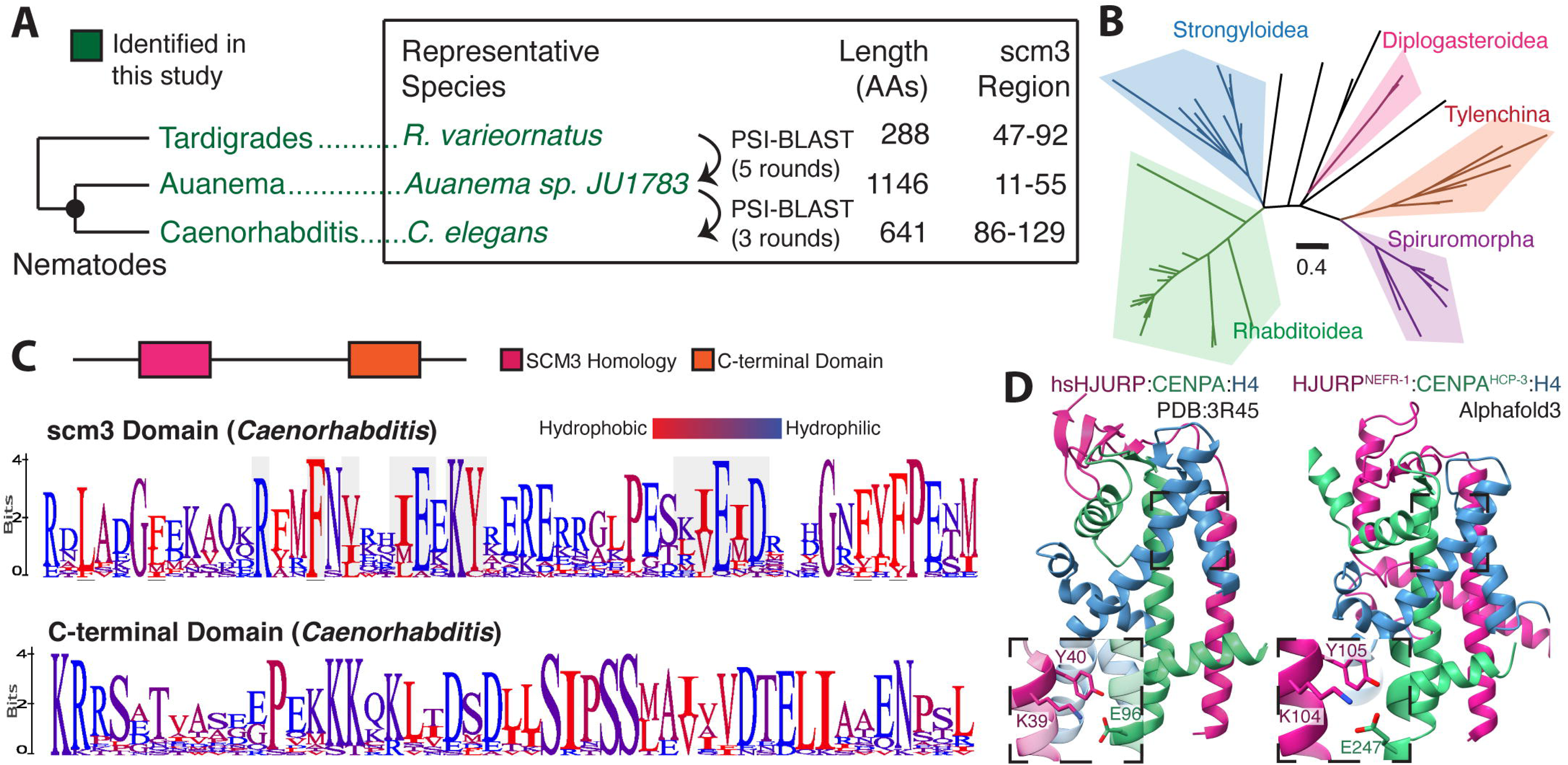
Nematode genomes encode a canonical HJURP protein. A. A visual schematic of the approach used to identify HJURP orthologs in nematodes. We use an intermediate *Auanema* predicted protein to establish scm3 homology in *Caenorhabditis*. The tardigrade scm3 domain provided the necessary signal for iterative searching of nematode proteomes. B. A phylogenetic tree constructed from scm3 domains of diverse nematode species is generally concordant with the species tree. C. *Caenorhabditis* HJURP proteins contain the N-terminal scm3 domain as well as a highly conserved C-terminal region. C-terminal logo plot shows a representative subsection of the entire conserved region. Alignments are from 20 species. D. A structural prediction of the *C. elegans* scm3 domain in complex with CENPA and H4 suggests structural features are broadly conserved with those of the human ortholog.

NEFR-1’s scm3-domain shares key conserved features with that of deuterostomes and fungi, including the KY motif, alternating (E/D) and (I/L/V) residues downstream of the KY, and a repeating hydrophobic pattern indicative of a helical region that could bind CENPA^HCP-3^ (Fig. 4C). This scm3-like region is broadly shared across *Caenorhabditis* despite little homology flanking it (Fig. S8B). There is also a conserved C-terminal motif shared across the genus, a feature highly reminiscent of the conserved C-terminal domain in *Drosophila* CAL1, which mediates its binding to CENPC. This motif may mediate a similar molecular interaction in *Caenorhabditis*. We used AlphaFold-multimer(*40*) to generate a predicted structure of the NEFR-1 scm3 domain in complex with CENPA^HCP-3^ and H4. This prediction shares a very similar global architecture to the human HJURP/CENPA/H4 structure, as well as a conserved interaction of the KY dipeptide and broad helix-sheet interface (Fig. 4D), strongly supporting our identification of NEFR-1 as a *bona fide* HJURP protein.

### C. *elegans nefr-1* encodes a *bona fide* HJURP ortholog

*C. elegans nefr-1* has been previously studied and described as coding for a neurofilament-like protein involved in neuronal function(*71*). Thus, our remote homology finding of an scm3 domain in NEFR-1 was initially puzzling. However, in addition to its remote homology and strict conservation in nematodes, including other *Caenorhabditis* species, we identified several attributes that bolster our hypothesis that *nefr-1* plays an important role in centromeric maintenance, a role that may have been overlooked in previous studies. For example, *nefr-1* expression is strongly correlated with genes that perform cell-cycle functions, as would be expected for centromeric function but not necessarily for neuronal function. The *nefr-1* gene is encoded in an operon with the DNA repair gene *msh-6* and shares a transcription factor-binding region with the *C. elegans* separase homolog *sep-1*, both of which are chromatin-associated, cell-cycle-regulated genes. Moreover, single-cell RNA-seq experiments reveal abundant *nefr-1* mRNA expression in the germline, as with CENPA^HCP-3^ (*73*, *74*), but not in neurons (Fig. S8C-F).

Buoyed by our success at validating drHJURP, we investigated the centromeric function of *nefr-1* in *C. elegans*. To date, no complete knockout for *nefr-1* has been functionally characterized (Fig. S9A). Previous analysis of *nefr-1* relied on an in-frame deletion allele, *ok471* (*C. elegans* Gene Knockout Consortium), which is viable. However, this mutant retains the putative scm3 domain and potentially the C-terminal conserved region, suggesting the possibility that several *nefr-1* function(s) may remain intact in the *ok471* truncation mutant. We used CRISPR-Cas9 to generate a *C. elegans* strain with an N-terminal stop codon and frameshift in the *nefr-1* gene (K15*), providing a near-complete *nefr-1* knockout model (Fig. S9A). Unlike the previously characterized deletion allele *ok471*, we were unable to isolate viable *nefr-1*(*K15\**) homozygotes, suggesting this ‘complete knockout’ mutation confers lethality or sterility. We therefore used a phenotypically marked balancer chromosome to follow the *nefr-1*(*K15\**) allele (Fig. S9B). While heterozygotes progressed with no discernible developmental delay and produced viable progeny, *nefr-1*(*K15*)* homozygous mutants predominantly arrested development at the L1 stage, were severely uncoordinated, and mostly died within 72 hours of hatching (Fig. S9C-D), all consistent with a previously reported aneuploidy phenotype(*75*). These results show that *nefr-1* is an essential gene.

We were initially puzzled by the ability of K15* homozygotes to progress to the L1 developmental stage, because failure to deposit CENPA^HCP-3^ should cause lethal defects in cell division even earlier during embryogenesis(*76*, *77*). We hypothesized that maternally deposited *nefr-1* mRNA or protein would be sufficient to enable sufficient CENPA^HCP-3^ deposition for faithful mitosis during embryogenesis and delay embryonic lethality. To unambiguously test NEFR-1 function without the confounding effects of maternal deposition, we generated an *nefr-1* strain with an N-terminal auxin-dependent degron tag (*AID::nefr-1*)(*78*, *79*) and used germline-specific expression of the auxin-dependent ubiquitin ligase TIR1 to achieve acute germline-specific depletion of NEFR-1. Embryos produced following NEFR-1 depletion from the hermaphrodite germline almost invariantly failed to hatch (Fig. 5A). These findings confirm our hypothesis that maternal deposition of *nefr-1* likely delays its embryonic lethality. To better understand the cause of lethality, we fixed and imaged early embryos after 24 hours of auxin-mediated NEFR-1 depletion. Whereas control embryos showed healthy mitotic features such as a uniform metaphase plate and homogeneous nuclei, auxin-exposed embryos displayed disorganized, highly variable chromosomes that lacked CENPA^HCP-3^ (Fig. 5B). These observations are consistent with a kinetochore-null phenotype(*76*), leading to severe chromosome segregation defects. We conclude that maternally contributed NEFR-1 is essential for embryonic cell division and deposition of CENPA^HCP-3^.

**Figure 5.**
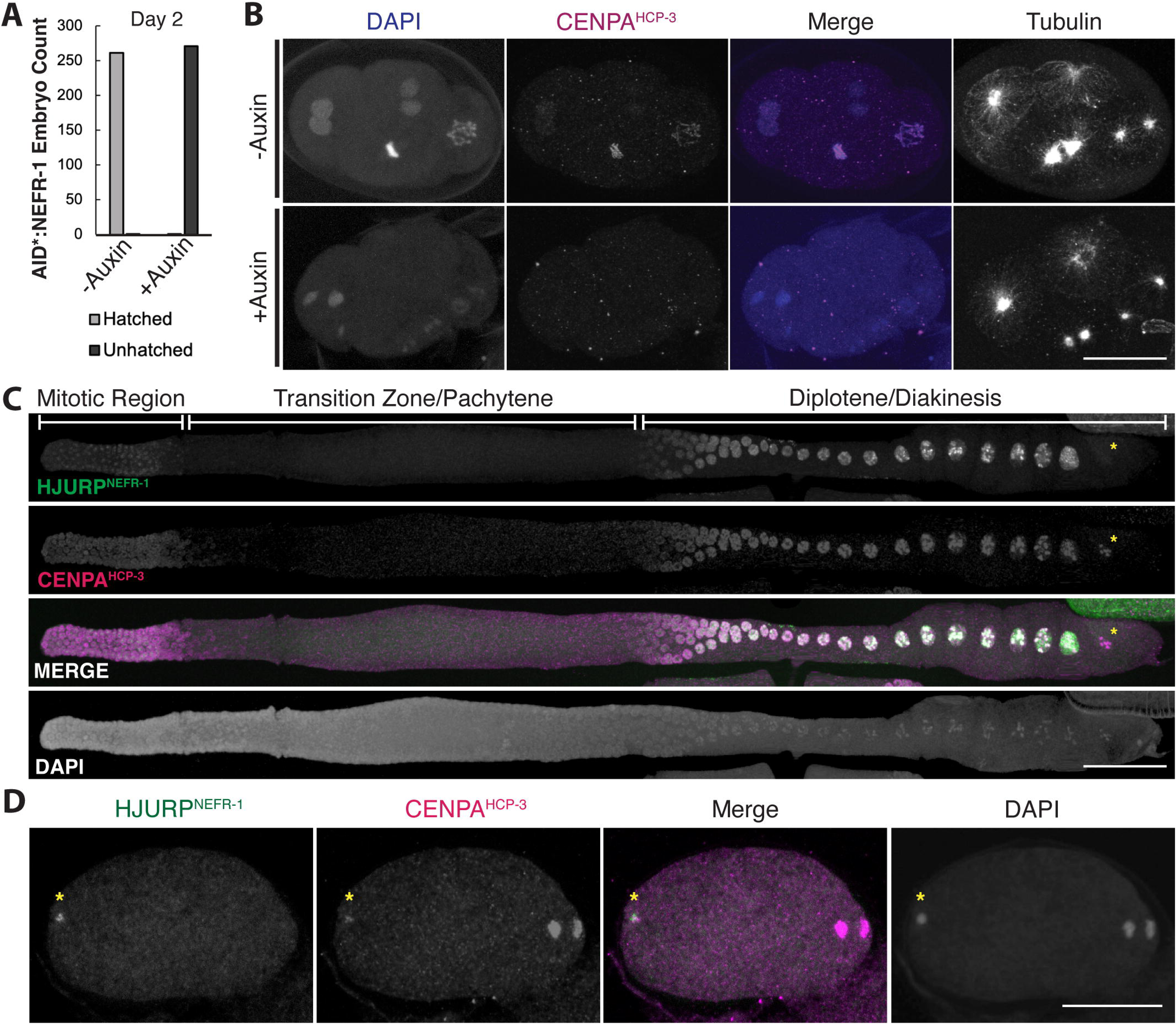
*C. elegans* NEFR-1 is an essential cell division regulator. A. Embryonic hatching counts from control or auxin-treated worms with an AID-tagged NEFR-1 show that NEFR-1 is essential for early embryonic development. B. Representative fixed imaging of control (top row) or auxin-treated (bottom row) 4-cell embryos with an AID-tagged NEFR-1. While control embryos have consistently sized, CENPA-coated chromatin, drug-treated embryos have catastrophic chromosome segregation abnormalities and lack CENPA on their chromatin. Scale bar is 20μm. C. NEFR-1 co-localizes with CENPA in the *C. elegans* germline, including at the mitotic distal tip (left) and diplotene/diakinesis (right). Yellow asterisk indicates that NEFR-1 leaves chromatin by late diplotene, while CENPA is retained for meiotic divisions. Scale bar is 40μm. D. A representative fertilized *C. elegans* oocyte undergoing anaphase I. NEFR-1 co-localizes with the paternal genome (yellow asterisk) but is absent from the maternal dividing chromatin (right), suggesting that the maternal machinery reloads CENPA onto the paternal DNA prior to the first zygotic division. Scale bar is 20μm.

Previous studies have shown that CENPA^HCP-3^ is loaded onto chromosomes *de novo* during the diplotene phase of *C. elegans* meiosis(*80*). To observe NEFR-1 localization, we generated an endogenous N-terminal 3xFLAG-tagged *nefr-1* allele (*3xFLAG::nefr-1*) and performed imaging in fixed *C. elegans* animals. We observed strong colocalization between CENPA^HCP-3^ and 3xFLAG::NEFR-1 in the germline, most prominently at the mitotic distal tip and on diplotene and diakinesis chromosomes, consistent with a centromere-defining role for NEFR-1 (Fig. 5C). We further observed that NEFR-1 localized to the paternal pronucleus within anaphase oocytes (Fig. 5D). This finding suggested a mechanism by which the paternal genome in *C. elegans*, which is devoid of CENPA^HCP-3^ in the spermatocyte(*80*), can re-establish centromeric chromatin prior to zygotic cell divisions. Like orthologous HJURP proteins(*6*, *7*, *56*, *81*), NEFR-1 appears to be temporally restricted, being absent from late prophase (Fig. 5C) and anaphase (Fig. 5D) meiotic chromosomes.

Finally, we performed immunoprecipitation-mass spectrometry (IP-MS) analysis using 3xFLAG::NEFR-1 from worm embryo lysate as bait to identify NEFR-1’s *in vivo* interaction partners. We found a unique association with CENPA^HCP-3^, as well as increased associations with cell cycle proteins, relative to untagged control lysates (Fig. S10A-B). As with zebrafish HJURP, we performed pulldown experiments using purified recombinant proteins and found a direct interaction between NEFR-1 and HCP-3/H4 that was stronger than that between NEFR-1 and H3/H4 (Fig. S10C). Together, these data confirm that *nefr-1* encodes the long-missing *C. elegans* HJURP ortholog that chaperones CENPA^HCP-3^ to chromatin and defines the holokinetic centromere in *C. elegans*.

### Scm3/HJURP orthologs are broadly conserved across *Metazoa*

Finally, we reinvestigated the early ancestry of the scm3 domain in *Metazoa*. In addition to arthropods, we extended our combination of PSI-BLAST and HMM-based searching with HHpred(*49*) validation (Fig. S3) to identify representative scm3 domains in the remainder of the protostome lineage as well as in the early-branching sponges and placozoa (Fig. 6A). We successfully identified Scm3/HJURP orthologs in most metazoan lineages we investigated (Fig. 6A), including in multiple species in which they were previously not found (Fig. S2), entirely filling the evolutionary gap between choanoflagellate Scm3 and human HJURP orthology first established 15 years ago(*9*) (Fig. S11). As in arthropods and deuterostomes, these sequences were highly variable in length and showed very little homology outside the scm3 domain. Structural prediction supports a unified scm3 protein fold across these early-diverging clades (Fig. 6B).

**Figure 6.**
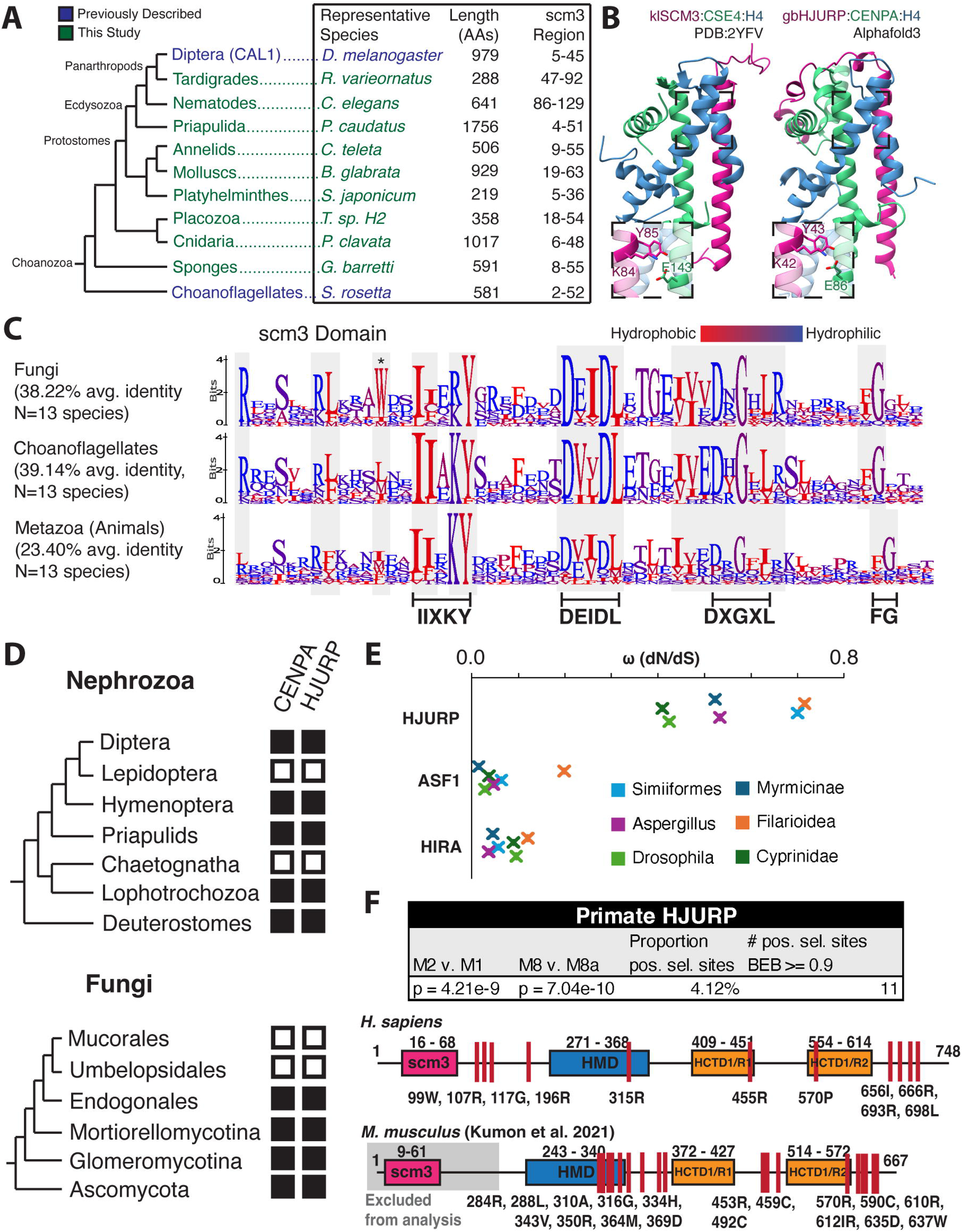
HJURP is present in basal metazoa. **A.** An expanded metazoan phylogeny encompassing basally-braching groups and protostomes for which we identified HJURP candidates. **B.** AlphaFold3 predicts a complex of the sponge *G. barretti* candidate HJURP with gbCENPA and H4, which is structurally similar to the published yeast *K. lactis* complex. **C.** Logo plots of representative Fungal, Choanoflagellate, and metazoan scm3 domains show select sequence features are conserved across opisthokont evolution. Metazoan scm3 domains show lower identity compared to those of the opisthokont sister clades. **D.** The loss of CENPA is associated with the absence of candidate HJURP proteins across opisthokont evolution. We failed to identify strong HJURP candidates in clades nested between those with clear sister group and outgroup HJURP orthologs. We show only a subset of clades with CENPA losses. **E.** HJURP evolves more rapidly than functionally comparable histone chaperones. Omega (dN/dS) values for HJURP candidates from well-sequenced clades were consistently higher than those for the histone chaperones ASF1 and HIRA. **F.** We identified sites of positive (diversifying) selection in primate HJURP in regions similar to (C-terminus, HMD) those previously identified in rodents (*90*), indicative of pervasive adaptive pressure.

Successful identification of scm3 domains in these lineages allowed us to investigate whether any features are unique to metazoan HJURP proteins relative to Scm3/HJURP proteins from yeast and choanoflagellates. We generated scm3 domain alignments encompassing major clades of fungi(*82*), choanoflagellates(*83*), and animals(*84*), respectively, of comparable species divergence timescales. We found that metazoan scm3 domains have lower overall sequence identity (Fig. 6C). Signatures such as the DXGXL motif, which were likely present in the common ancestor of fungi and animals, appear to evolve under less stringent selective constraints on the metazoan branch. We also found that the acidic/short hydrophobic region immediately following the KY, which interacts specifically with the CENPA CATD beta-strand, faces relaxed constraints in metazoans relative to other opisthokonts (Fig. 6C). One reason for this rapid divergence might be the evolutionary acquisition of asymmetric female meiosis in *Metazoa*, which may have led to the rapid evolution of CENPA proteins (*85*, *86*) and, subsequently, of HJURP proteins.

Our analyses indicate that some unusual features of CENPA chaperones and their presence have independently converged in yeast and metazoans. For example, *Dikarya* (a subkingdom of *Fungi*) largely prefer a tryptophan in the same position as CAL1 in *Diptera* (asterisk, Fig. 6C). In both *Fungi* and *Metazoa*, the loss of CENPA also coincided with the loss of HJURP; such dual losses have occurred multiple times across the opisthokonts (Fig. 6D) (*87–89*). This is a further indication that HJURP and CENPA are functionally co-dependent, supporting these candidate proteins as *bona fide* functional HJURP orthologs.

Previous work found evidence of positive selection on rodent HJURP (*90*) and mosquito CAL1(*35*) orthologs. Given their extreme divergence across clades, we conducted dN/dS analyses of candidate HJURP genes from selected taxa, each of which contains several genomes, to test for rapid evolution. Indeed we found that high nonsynonymous substitution rates are a general feature of this gene: dN/dS values were on average an order of magnitude higher for HJURP genes than for the functionally similar histone chaperones HIRA and ASF1 (*91*) in multiple metazoan lineages, as well as in fungi (Fig. 6E). Additionally, we found evidence of site-specific positive selection across primate evolution, reminiscent of previous observations in rodents (*90*), identifying HJURP as a gene under constant innovation across long evolutionary timescales (Fig. 6F). Much like HJURP in *Murinae*, these regions of selection were concentrated in the C-terminus and around the HMD region, suggesting persistent selective pressure across millions of years of evolution and implicating a functional role in these uncharacterized regions. This remarkably high nonsynonymous substitution rate may have accelerated HJURP’s sequence divergence across several metazoan lineages, necessitating the use of remote homology detection methods employed in this study. Consistent with the scm3 domain as the most conserved functional domain, we did not identify residues under positive selection in the N-terminal region of primate HJURP.

## Discussion

HJURP^Scm3^ is a fundamental component of the centromere identity-defining machinery, essential for sustained error-free cell division(*1*). Despite its cardinal function, HJURP has a paradoxically rapid evolutionary rate – so rapid that it was believed to be missing from many animal genomes(*29*). By combining PSI-BLAST, HMM-based searches, AlphaFold-multimer predictions, and functional genetic and cytological experiments, we significantly revise this model by demonstrating that *bona fide* Scm3/HJURP orthologs are in fact broadly retained across *Metazoa* but have been obscured by extreme sequence divergence. Our work on HJURP^Scm3^ supports the growing evidence that “losses” of essential genes more often reflect failures of homology detection(*92–94*) and proposes a structure-informed remote homology detection pipeline to uncover such “missing” genes (Fig. S3).

Recent advances in sequencing and structure prediction have made challenges in remote homology detection appreciably more tractable. However, remote homology detection efforts are still often aided by evolutionary intermediates that act as ‘stepping stones’, providing the necessary signal to connect distinct clades. Indeed, the choanoflagellate HJURP served as an intermediate, providing the link to establish orthology between yeast Scm3 and mammalian HJURP proteins. In this study, we found that genes from contentious “living fossil” genomes (*95*), an informal collection of slowly evolving genomes, such as those of the coelacanth and horseshoe crab, provided similar links for identifying or confirming HJURP orthologs. Even when the genes are unannotated, as is the case for coelacanth, horseshoe crab, *Hymenoptera*, or the nematode *Auanema,* they can provide the ‘stepping stone’ to establish homology. Intriguingly, the *L. chalumnae* hit initially served as such a critical intermediate despite itself lacking an intact scm3 domain.

HJURP is composed predominantly of polar residues and disordered domains, making it a poor candidate for purely structure-based searching. The combination of structure-informed alignments with HHpred and predictions with Alphafold-Multimer, however, greatly reduced the false discovery rate. Continued development of *in silico* screening methods that search for oligomer-specific conformations, such as that of HJURP bound to the CENPA/H4 dimer, will exponentially increase the throughput of this type of remote homology detection.

The use of model organisms to study the cell cycle has been paramount to understanding how and when cell-division perturbations manifest during development, how tissues idiosyncratically employ cell-cycle differences, and how germline tissues fundamentally differ from the soma. By identifying *bona fide* HJURP orthologs in the genetically tractable zebrafish and *C. elegans* models, our work opens avenues for understanding how centromeric identity differentially affects processes in multicellular organisms. For example, our work suggests that the *C. elegans* paternal genome, which lacks CENPA(*80*) in spermatocytes, reacquires CENPA in the oocyte from maternal machinery prior to the first zygotic division. A broader toolset of model organisms with more complete repertoires of centromeric protein orthologs will allow both rapid and mechanistic dissection of core centromeric functions, while also providing fodder for comparative analyses of the plasticity and evolvability of centromeric apparati across organisms.

Although HJURP sequences are predominantly diverged, the functional principles that govern HJURP are broadly conserved. Across distant lineages, HJURP orthologs preferentially bind CENPA–H4 over H3–H4, localize to centromeres in a cell-cycle-dependent manner, colocalize with CENPA, and are essential for chromosome segregation and development. AlphaFold-Multimer shows that HJURP–CENPA–H4 complexes from fungi, zebrafish, humans, nematodes, and insects adopt strikingly similar three-dimensional folds, with conserved hydrophobic pockets and KY-centered motifs that maintain recognition of their respective CENPA proteins via their CATD domains despite minimal primary-sequence identity at contact residues. Other features, such as CENPC-binding(*24*, *36*, *54*) and DNA-binding(*96–100*), also appear to be conserved over long evolutionary timescales.

Despite this broad conservation, lineage-specific adaptations can also be found in HJURP proteins. For example, HJURP proteins from vertebrates, tetrapods, and teleost fish share additional motifs immediately upstream of the DNA-binding HMD region and within the Mis18-binding R2 region that are absent in echinoderms. Perhaps the most striking lineage-specific adaptations can be found in insect CAL1 proteins. Although the scm3 domain is the most conserved region across metazoans, it shows relatively low homology in *Diptera* compared with other arthropods. In contrast, a downstream region immediately C-terminal to the scm3 domain exhibits higher sequence conservation. Furthermore, Diptera have evolved a structurally distinct CENPC-binding domain at the C-termini of their CAL1 proteins (*36*). This finding, combined with the variable and incompletely defined contribution of CENPC to HJURP recruitment, even among tetrapods(*23*, *24*, *101*), exemplifies how centromeric machinery can be substantially rewired while maintaining functional fidelity.

HJURP genes evolve roughly an order of magnitude faster than other histone chaperones, such as HIRA and ASF1, in *Metazoa*. The previous inability to correctly identify HJURP orthologs in some metazoans is almost certainly a consequence of this more rapid evolution. Although the specific selective pressures driving HJURP’s diversification are still unknown, the accelerated rate of divergence we observe in metazoan scm3 domain-containing proteins, combined with their demonstrated direct role in modulating centromere identity, makes it very likely that these candidate HJURP proteins play an important role in suppressing “selfish” centromeric DNA during ‘centromere drive’(*29*, *85*, *102*). The centromere drive model posits that centromeric DNA variants bias their own transmission during asymmetric female meiosis, during which only one of four meiotic products is retained in the oocyte. Conversely, DNA-binding centromeric proteins, such as CENPA(*86*), diversify to restrict the biased transmission of “selfish centromeres” during female meiosis in both plants and animals.

We hypothesize that HJURP may evolve under dual pressures from centromere drive. First, HJURP’s distinct DNA-binding regions(*97*, *99*, *103*) might adapt to centromeric DNA changes to effectively modulate centromere identity. Second, HJURP must keep pace with CENPA’s rapidly evolving CATD region(*65*) to ensure high fidelity of its preferential interaction with CENPA/H4 over H3/H4. Consistent with this prediction, as recently described in rodents(*90*), we observe strong signatures of positive selection in primate HJURP, suggesting its role in mitigating this persistent genetic conflict. However, we and others(*104*) also identify rapid evolution in fungal Scm3, which is not thought to face the same selective pressures from centromere drive as animal HJURP(*105*), as fungi typically undergo only symmetric meiosis. Our findings suggest that evolutionary forces both during and beyond female meiosis drive the adaptive evolution of Scm3/HJURP chaperones. Our pipeline for identifying extremely remote homology enables an understanding of the essential molecular players in these ancient genetic conflicts and provides a means to connect distant homologies in centromeric biology and beyond.

## Materials and Methods

### HJURP Candidate Searching and Selection Criteria

A schematic representation of the approach is shown in Fig. S3. Initial HJURP candidates were identified with a combination of iterative BLAST searching (PSI-BLAST, expected threshold 5 and PSI-BLAST threshold 0.5)(*43*) and Hidden Markov Model searching (HMMsearch, default settings)(*63*, *64*). Proteomes searched include non-redundant protein sequences from NCBI, the UniProtKB/TrEMBL complete database (https://www.ebi.ac.uk/training/online/courses/uniprot-quick-tour/the-uniprot-databases/uniprotkb/), Wormbase(*106*), and InsectBase(*107*). HMMs were generated from MUSCLE(*45*) alignments (using default settings) of representative tetrapod, Dipteran, and/or choanoflagellate/yeast sequences. Queried proteomes were searched via the Fred Hutchinson computational cluster implementation of HMMer (using default settings)(*63*). Initial HMM hits were not limited by any statistical cutoff, although common false-positive signals (including from an insect E4 esterase-like gene) were excluded; the intentional omission of a strict statistical cutoff allowed us to obtain a more holistic interpretation of remote homology data. Anecdotally, we found that strong hits were often within largely unstructured, polar protein sequences, although we did not formally use this criterion for filtering.

Following initial queries, sequences were analyzed in HHpred(*49*) to confirm regions structurally and sequentially homologous to published scm3 protein domains (PDBs 2L5A(*10*), 2YFV(*11*), 3R45(*12*), 6XWV and 6XWS(*36*)). HHpred E-values ranged widely (Table S1) but were generally below 1. These hits were subsequently used for trimeric AlphaFold-Multimer (using default setting)(*41*) predictions of the candidate scm3 domain with a single copy each of the respective species’ canonical H4 and CENPA. Only annotated sequences with positive HHpred (default settings)(*49*) scm3 predictions, successful scm3-like structure predictions, and reciprocal PSI-BLAST or HMMer searches to published Scm3/HJURPs were used for downstream processing. HMMs were updated to include these new sequences, allowing for iterative searching.

The following candidate proteins are presented in the main text figures. **Figure 1**: *L. chalumnae* (XP_006012693.3, PSI-BLAST E-value 6e-04), *D. rerio* (NP_001076347.1, PSI-BLAST E-value 2e-06), *S. torazame* (GCB61218.1, PSI-BLAST E-value 5e-12), *P. marinus* (XP_032806972.1, PSI-BLAST E-value 0.2), *B. belcheri* (XP_019630933.1, PSI-BLAST E-value 0.03), *P. miniata* (XP_038057064.1, PSI-BLAST E-value 0.77). **Figure 3**: *N. vitripennis* (XP_001603895.3, HMMer E-value 0.99), *G. longicerus* (A0AAN9VWU2, HMMer E-value 2.7e-04), *D. pulex* (EFX88527.1, HMMer E-value 1.9e-06) *A. ventricosus* (GBM08647.1, HMMer E-value 0.46), *L. polyphemus* (XP_022237972.1, HMMer E-value 3.3e-03), *R. varieornatus* (GAU99461.1, HMMer E-value 1.3e-12). **Figure 4**: *Auanema sp. JU1783* (CAI4225445.1, PSI-BLAST E-value 0.44), *C. elegans* (NP_491161.2, PSI-BLAST E-value 7.0e-19). **Figure 6**: *P. caudatus* (XP_014675948.1, PSI-BLAST E-value 6.0e-03), *C. teleta* (ELT94853.1, PSI-BLAST E-value 4.0e-03), *B. glabrata* (KAI8798300.1, HMMer E-value 3.9e-15), *S. japonicum* (TNN07270.1, HMMer E-value 1.8e-12) *T. sp. H2* (RDD41302.1, HMMer E-value 1.9e-16), *P. clavata* (CAB3982230.1, HMMer E-value 2.0e-05), *G. barretti* (CAI8037030.1, HMMer E-value 3.5e-07).

A table of representative HJURP candidate proteins across Metazoa is provided in Table S1, and HMM model alignments are in Data S1. Per-residue AlphaFold pLDDT (confidence) scores are mapped onto predicted structures in Supplemental Figure 12.

### Structure Analyses

RMSD backbone calculations and structure alignments, contact determination, and visualizations were performed in UCSF ChimeraX-1.10(*108*) using the matchmaker tool.

### Sequence Alignments

Protein sequence alignments were generated with the MUSCLE algorithm (default settings) in Geneious. For HMM model building and scm3 domain visualization, alignments were extensively manually trimmed to remove species-specific insertions. Logo plots(*109*) were generated from trimmed alignments and colored by hydrophobicity in Geneious. Frequently, the only region successfully aligned across order-level alignments was the N-terminal scm3 region, and it was often the only region used for further analyses. Protein alignments are available in Data S2.

Codon alignments were performed using the webPRANK(*110*) server on whole-gene coding-sequence data from NCBI and were subsequently trimmed manually. Sequences containing large insertions or with exceptionally long phylogenetic branches were excluded from final analyses. Only clades with at least 8 sequences were used for alignments and analysis of evolutionary selective pressures. Codon alignments are available in Data S3.

### Phylogenetic and dN/dS Analyses

All phylogenetic trees were generated via the IQ-Tree(*111*, *112*) web server, allowing for free-rate heterogeneity and using optimized substitution models, and then visualized in FigTree v1.4.4 (https://github.com/rambaut/figtree). The phylogenies presented in Figures 1 and 4 were generated from the scm3 domain alignment alone, whereas the tree in Figure 3 was generated from whole-gene alignments.

Whole-gene dN/dS calculations and site-specific selection tests were performed using maximum-likelihood estimation with the codeml algorithm in PAML (*113*). Model 0 was used to calculate whole-gene dN/dS. Site-specific positive selection was calculated as previously described(*114*) for primate HJURP. Briefly, models that either include (models 2, 8) or exclude (model 1, 8a) a category allowing dN/dS> 1, indicating positive selection, were compared. Codons with Bayes Empirical Bayes scores above 0.9 for model 8 were highlighted as evolving under positive selection.

### Recombinant Protein Expression and Purification

For MBP-drHJURP200 expression, the first 200 amino acids of zebrafish HJURP were amplified from 2dpf cDNA and cloned into an expression vector with an N-terminal MBP tag and C-terminal 6XHIS tag. Protein expression and purification were carried out largely as in(*53*) with slight modifications. Briefly, BL21 DE3 LOBSTR cells were transformed with the drHJURP expression vector and grown in Luria Broth with ampicillin at 37°C to an optical density (OD) of 0.6, after which they were induced with 1mM of IPTG and shifted to 18°C with shaking for overnight protein expression. The following day, cells were resuspended in HisTrap binding buffer (20mM Na_3_PO_4_, 500mM NaCl, 20mM imidazole pH 7.4) with a cOmplete protease inhibitor tablet (Roche) and lysed via sonication. Following clarification at 30000 × g for 20 minutes at 4°C, the soluble lysate was run over a 5mL HisTrap column equilibrated in binding buffer. The column was washed with 50mL of binding buffer, and protein was eluted with HisTrap elution buffer (20mM Na_3_PO_4_, 500mM NaCl, 500mM imidazole, pH 7.4). The eluate was then further purified via preparative size exclusion chromatography on a Superdex 200 Increase 10/300 column (Cytiva) in a buffer with 25mM Tris-HCl pH 7.5, 300mM NaCl, 5mM β-mercaptoethanol. Fractions containing pure protein in 10% v/v glycerol were snap-frozen in liquid nitrogen and stored at –80°C. A similar approach was used to prepare the NEFR-1 construct, which consisted of amino acids 68-196 of *C. elegans* NEFR-1, except that the HisTrap eluate was further purified using a Superdex 75 Increase 10/300 column.

Soluble drCENPA/H4 tetramers were purified as an adaptation of Servin and Straight(*115*). drCENPA was amplified from 2dpf zebrafish embryos and cloned into a bicistronic vector encoding both drCENPA and H4. The vector was transformed into BL21 DE3 cells, grown to an OD of 0.6 at 37°C, then induced to express the tetramer with 1mM IPTG. Following an additional 5 hours of growth, cells were harvested and lysed in histone binding buffer (20mM Tris-HCl pH 7.4, 750mM NaCal, 10mM β-mercaptoethanol, 0.5mM EDTA) with a cOmplete protease inhibitor tablet (Roche). The lysate was cleared at 30000 × g for 20 minutes at 4°C, then bound to a HiTrap SP FastFlow column and washed with binding buffer. The tetramer was eluted with a 20-column-volume gradient into binding buffer containing 2 M NaCl. Fractions containing tetramer were further purified via preparative size exclusion chromatography on a Superdex 200 Increase 10/300 column (Cytiva) in histone binding buffer. Fractions containing pure tetramer were pooled, concentrated, snap-frozen in 10% v/v glycerol, before being stored at –80°C. A similar approach was used to prepare soluble HCP-3/H4 tetramers, except that a truncated form of HCP-3, residues 148-288, was used. Soluble H3/H4 tetramers were purchased commercially (New England Biolabs #M2509).

### Amylose Pulldown

Amylose resin pulldowns were performed as in Hori et al.(*54*) with amylose resin replacing GST resin. 50pmol of MBP-drHJURP or MBP-NEFR-1 was incubated with 30μL of amylose resin equilibrated in pulldown buffer (20 mM HEPES-NaOH, pH 7.4, 750mM NaCl, 0.05% NP-40, 5mM DTT) for 1 hour at 4°C. The resin was then washed three times in pulldown buffer and incubated with 25 pmol of drCENPA/H4, HCP-3/H4, or H3/H4 tetramer, as indicated. The resin was washed three more times in pulldown buffer, then proteins were eluted in 30μL of SDS-PAGE buffer and analyzed via SDS-PAGE stained with GelCode Blue.

### Zebrafish Care and Maintenance

*Danio rerio* animals were maintained at the Fred Hutch Cancer Center in compliance with IACUC-approved protocols, and experiments were conducted in accordance with IACUC standards. Zebrafish were bred and maintained according to an established protocol(*116*). Sex is not a relevant variable in our experiments, which were performed before the onset of sex determination in zebrafish(*117*). No transgenic zebrafish lines were used.

### Zebrafish CRISPant Generation

CRISPant(*118*) mutant alleles were generated using CRISPR/Cas9 with standard protocols (*119*). gRNAs were designed using chopchop (https://chopchop.cbu.uib.no), ordered as Alt-R sgRNAs (IDT), and pooled in a 1:1 manner for each of three conditions: 1) Control guide(*118*) with two HJURP scramble guides, 2) three drHJURP guides, and 3) two drCENPA guides. One-cell-stage embryos were injected with 1nL of 10μM pooled gRNA and 5μM Cas9 protein (IDT, #1081058), and F0 mutant animals were sampled for editing efficiency via PCR. Guide RNAs and genotyping primers used in this study are provided in Table S2. Unfertilized embryos were removed after 2hpf, and CRISPant morphology was scored on 24hpf larvae by eye compared to uninjected control individuals. “Necrotic” individuals had wild-type morphology but noticeable darkened opaque tissue, while “morphological defects” were marked by differences in body shape compared to uninjected references.

### Zebrafish Fluorescence Microscopy

For colocalization experiments, drHJURP and drCENPA coding sequences were amplified from 2dpf larval cDNA and cloned into fluorescent tag backbones via standard cloning techniques. mRNAs were generated from template plasmid DNA via *in vitro* transcription (mMessage mMachine, Invitrogen) and purified via gel extraction. 100pg of each purified mRNA was injected into one-cell-stage embryos in a volume of 1nL(*120*). For CRISPant analyses, 1nL of 100ng/uL of H2B-mCherry mRNA was injected together with 5μM Cas9 protein and 10μM of pooled HJURP or control CRISPR guides. For CENPA intensity calculations, these injections were supplemented with 100 ng/uL eGFP-drHJURP. Construct sequences and amplification primers for fluorescent proteins are provided in Table S2.

6hpf individuals were embedded in 1.0% low-melt agarose and imaged on a Leica SP8 inverted confocal microscope using a 28°C heated stage and 1.5% laser power at 488nm and 555nm. Stacks with a 1μm step size were collected for 5 hours at 6-minute intervals. Representative still frames of different cell cycle stages are shown in Figure 2. Images were processed in FIJI(*121*). For anaphase fidelity quantification, anaphases from time-lapse images were scored as “erroneous” if they included lagging or tethered mCherry-H2B signal between segregating chromatin volumes at the second frame of anaphase onward, while all others were considered wildtype. For drCENPA intensity quantification, 10-image stacks from 10 hpf time-lapse images were max intensity projected, equally thresholded on eGFP intensity to remove background nuclear signal and only isolate centromeric signal, and measured as a ratio of eGFP-drHJURP:mH2B-mCherry per nucleus.

### C. *elegans* Maintenance and Genetics

*C. elegans* was maintained at 20°C on standard nematode growth media (NGM) and fed *E. coli* OP50, unless otherwise noted. Genotyping primers used in this study are provided in Table S2. A list of strains used in this study is provided in Table S3.

For *nefr-1* K15STOP embryo viability counts, genotypes were tracked via the presence of the balancer tmC20 which causes GFP expression in the pharynx of heterozygotes and a dumpy phenotype with GFP in homozygotes with wildtype *nefr-1*. Single heterozygotes were allowed to lay eggs overnight and then removed from the plate the following morning. Following a 48-hour hatching period, worms were genotyped and scored for developmental stage based on standard criteria. Each day following, worms were scored on both developmental stage and survival until 7 days post-laying.

For auxin hatching experiments, three NJL5706 L4 worms were placed on NGM + OP50 plates layered with 500μL 500μM K-NAA (l1-Naphthaleneacetic Acid Potassium Salt) in M9 medium or control plates with 500μL of M9 alone. The next afternoon, worms were transferred to fresh plates containing auxin or control solutions and allowed to lay eggs overnight. The next morning, adult worms were removed from the plates, and their eggs were allowed to hatch for 48 hours. Plates were scored on the number of eggs that hatched.

### C. *elegans* CRISPR/Cas9 Genome Modification

CRISPR/Cas9 genome editing was performed by microinjection of guide RNA/Cas9 ribonucleoprotein (RNP) complexes (IDT #1072532, #1081058) and custom crRNA along with either ssDNA oligos (IDT) or partially-melted dsDNA oligos as homology-directed repair (HDR) templates(*122*, *123*). See Table S2 for guide and repair constructs used for microinjection.

### C. *elegans* Fluorescence Microscopy

*C. elegans* immunofluorescence samples were prepared as described(*124*, *125*) with slight modifications. Gravid adults were dissected into a 15μL drop of M9 solution on charged slides (ProbeOn Plus, Fisher) and fixed with a 15μL drop of 2% PFA (Sigma) in M9 for 5 minutes. As much liquid as possible was wicked away with blotting paper. The drop was covered with a 1.5 coverslip, and the slide was frozen in liquid nitrogen for 5 minutes. The slide was then removed, the slip quickly cracked off with a razor blade, and the slip was dipped into ice-cold methanol for 1 minute. Slides were then synchronized in PBS-T (PBS with 0.1% Triton X-100). Following synchronization, slides were washed twice in PBS-T, then blocked with a 100μL drop of AbDil (PBS pH 7.4, 4% BSA, 0.1% Triton X-100, 0.02% NaN_3_) at room temperature for 1 hour. Samples were incubated overnight in 100μL of primary antibody diluted in AbDil at 4°C. The next day, slides were washed 3 times in PBS-T, then incubated with 100μL of secondary antibody for 1-2 hours at room temperature. Slides were washed 3 times in PBS-T and mounted with 8μL of DAPI-containing antifade media. The following antibodies were used: anti-FLAG M2 (1:500, Sigma F1804), anti-HA tag (1:250, Abcam ab236632), anti-αTubulin (1:250, Abcam ab7291), anti-Rabbit AlexaFluor 555 (1:500, Invitrogen A-21428), anti-Mouse AlexaFluor 488 (1:500, Invitrogen A-11001).

Exposure-standardized images from samples were collected on a Leica STELLARIS8 with Leica Application Suite X software. Stacks of approximately 30μm were collected with an oil objective at either 40X (germlines) or 63X (eggs/embryos). Images were processed in FIJI(*121*), in which color channels were separated, recolored, and merged; DAPI channels were scaled for presentational clarity, and germlines were computationally straightened. All images are maximum-intensity projections of z-stacks that encompass the entire germline or embryo.

### C. *elegans* NEFR-1 Immunoprecipitation

Immunoprecipitations from worm embryo lysate were performed as described(*126*) with slight modifications. Worms from mixed-stage populations were bleach-synchronized and hatched for 24 hours in liquid M9. Synchronized worms were then plated on 20X-concentrated OP50 for large-scale growth and allowed to progress to the gravid adult stage for 3 days. Worms were then bleached, washed, pelleted, and flash-frozen in liquid nitrogen for downstream processing.

Worm pellets were thawed on ice and resuspended 1:0.5 in 1.5X lysis buffer(*127*) (50mM HEPES pH 7.5, 1mM EGTA, 1mM MgCl_2_. 150mM KCl, 15% glycerol, 0.1% NP-40, 0.75mM DTT, cOmplete protease inhibitor tablet, 1X Roche PhosSTOP). Worm suspensions were lysed via sonication for fifteen 30s on/30s off cycles using a Bioruptor Pico sonication device (Diagenode). Lysates were cleared at 19000 × g for 20 minutes at 4°C. Clarified supernatants were transferred to fresh 1.5mL tubes and kept on ice for all subsequent steps. Samples were diluted in lysis buffer to 10mg/mL in 200 μL aliquots and incubated for 2 hours with 100 μL of lysis buffer-equilibrated magnetic anti-FLAG M2 beads (Sigma M8823). Following incubation, beads were isolated with a magnetic stand, washed 3X in wash buffer and 2X in 50mM HEPES pH 7.5, and boiled 1:1 in SDS-PAGE buffer for downstream mass spectrometry analyses.

Mass spectrometry data collection and processing were performed at the Fred Hutchinson Proteomics and Metabolomics core facility. Samples were desalted and run on an Orbitrap Ascend. The raw mass spec data were searched using Proteome Discoverer (PD) version 3.1 against a Uniprot C. elegans database, along with common contaminants. The search results were filtered for high confidence, 1% false discovery rate at the peptide level. The ratio of the sum PEP score was used to determine the relative representation of peptides within the 3xFLAG::NEFR-1 strain compared to an untagged control (Table S4). Peptides not found in the negative control were assigned a sum PEP score of 0.5 for comparison. All ratios were normalized to vitellogenins, the most common contaminant in both samples.

## Acknowledgments

We would like to thank members of the Malik and Campbell labs for valuable discussions and Sue Biggins, Pravrutha Raman, Peter Dietzen, and Maria Toro Moreno for their critical reading of the manuscript. *C. elegans* strain FAS131 was a gift from Florian Steiner.

## Author Contributions

Conceptualization: JAH, MGC, HSM

Methodology: JAH, JAS, IT, JMY, NJL, HSM

Software: JMY Validation: IT, HSM

Formal analysis: JAH, HSM

Investigation: JAH, JAS, IT, JMY, HSM

Resources: JAH, JAS, IT, CBM, NJL, MGC, HSM

Data curation: HSM

Writing—original draft: JAH, HSM

Writing— review & editing: JAH, JAS, IT, JMY, CBM, NJL, MGC, HSM Visualization: JAH, HSM

Supervision: JMY, CBM, NJL, MGC, HSM

Project administration: NJL, CBM, MGC, HSM

Funding acquisition: JAH, CBM, NJL, MGC, HSM

## Data and Materials Availability

All data needed to evaluate the conclusions in the paper are present in the paper and/or the Supplementary Materials.

## Table of contents for supplemental material

Figures S1 to S12

Tables S1 to S4

Data S1 to S3

Movie S1

