## Supplementary material for "Remote homology and functional genetics unmask deeply preserved Scm3/HJURP orthologs in metazoans": Fig S

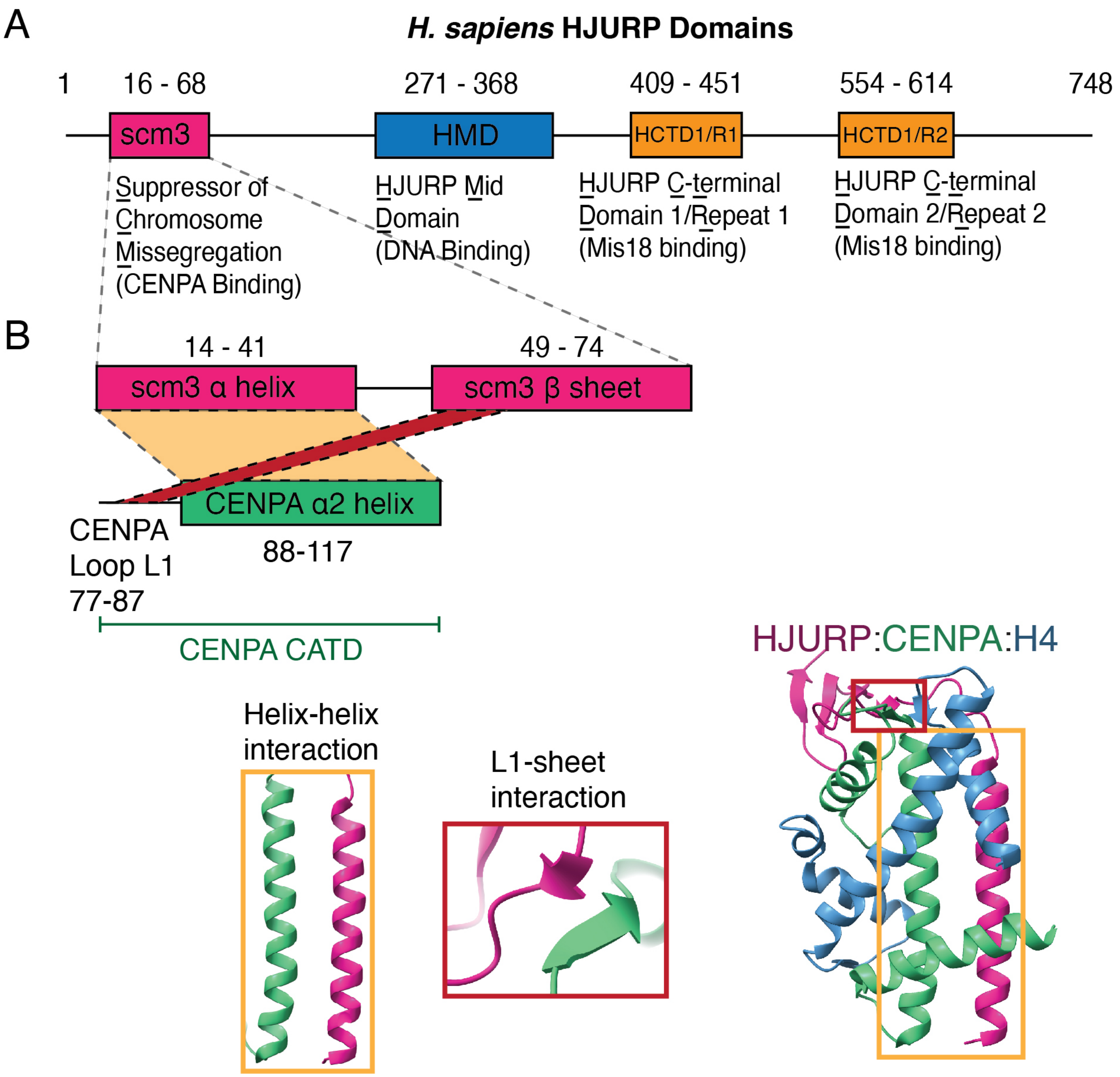


**Figure S1. Structure-function relationships of the human HJURP protein. A.** A domain schematic for human HJURP. The suppressor of chromosome missegregation (scm3) domain directly binds CENPA/H4 dimers, the HJURP mid domain (HMD) is implicated in DNA binding, and HJURP C-terminal domains (HCTD) 1 and 2 interact with Mis18 to recruit HJURP to the centromere. **B.** A structural overview of human HJURP-CENPA interactions. CENPA forms an antiparallel helix and strand with HJURP. Structure is PDB 3R45[12].

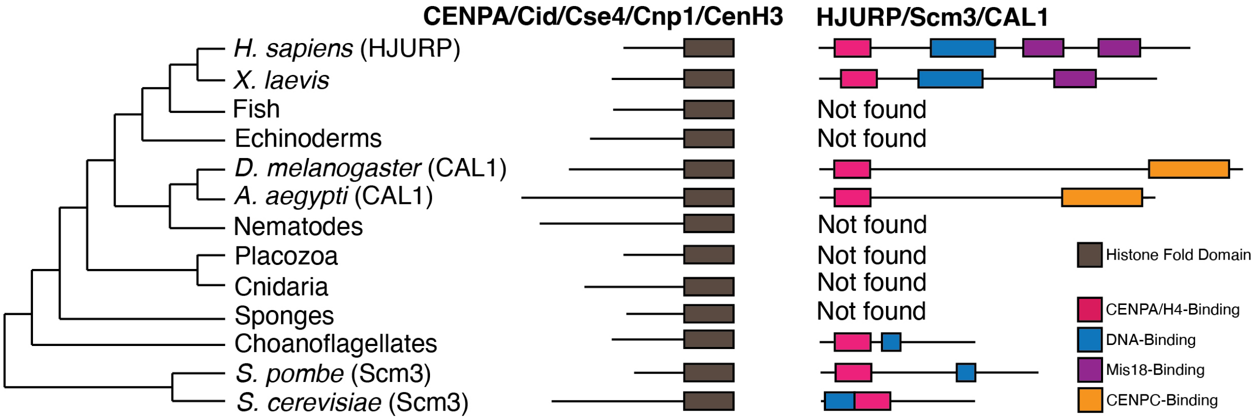


**Figure S2. HJURP orthologs had not previously been found in most metazoan clades. A.** An unscaled phylogeny showing relationships among major metazoan groups. Select scm3 domain-containing CENPA chaperones have been independently identified and biochemically characterized across the animal tree, yet many species with CENPA appear to lack a scm3 chaperone.

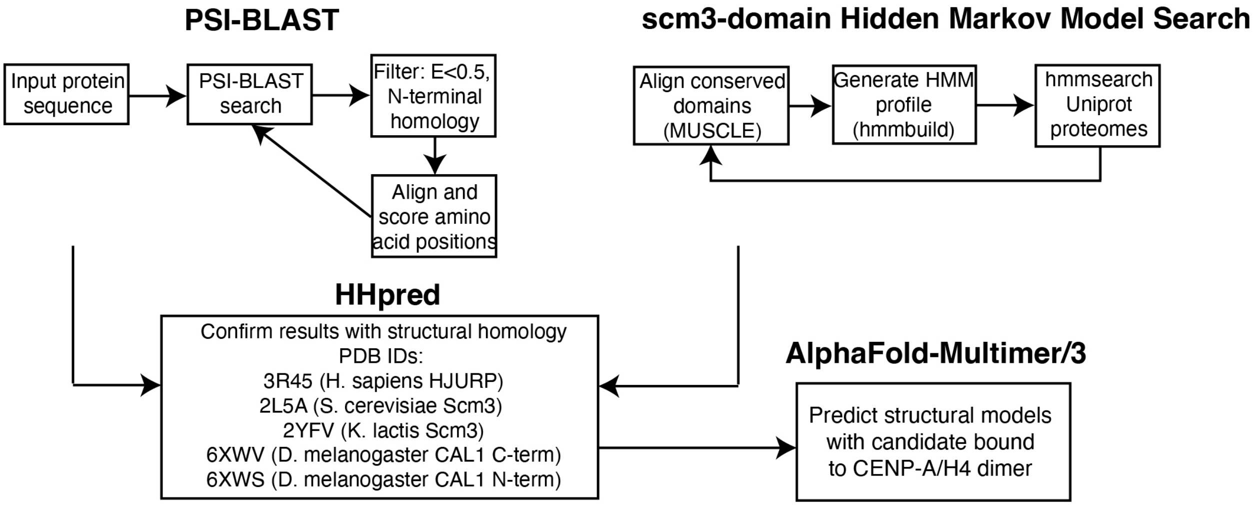


**Figure S3. A schematized approach for detecting remote HJURP orthologs.** We employed a holistic homology search pipeline integrating sequence and structural data to identify and confirm distant Scm3/HJURP orthologs that previously escaped detection.

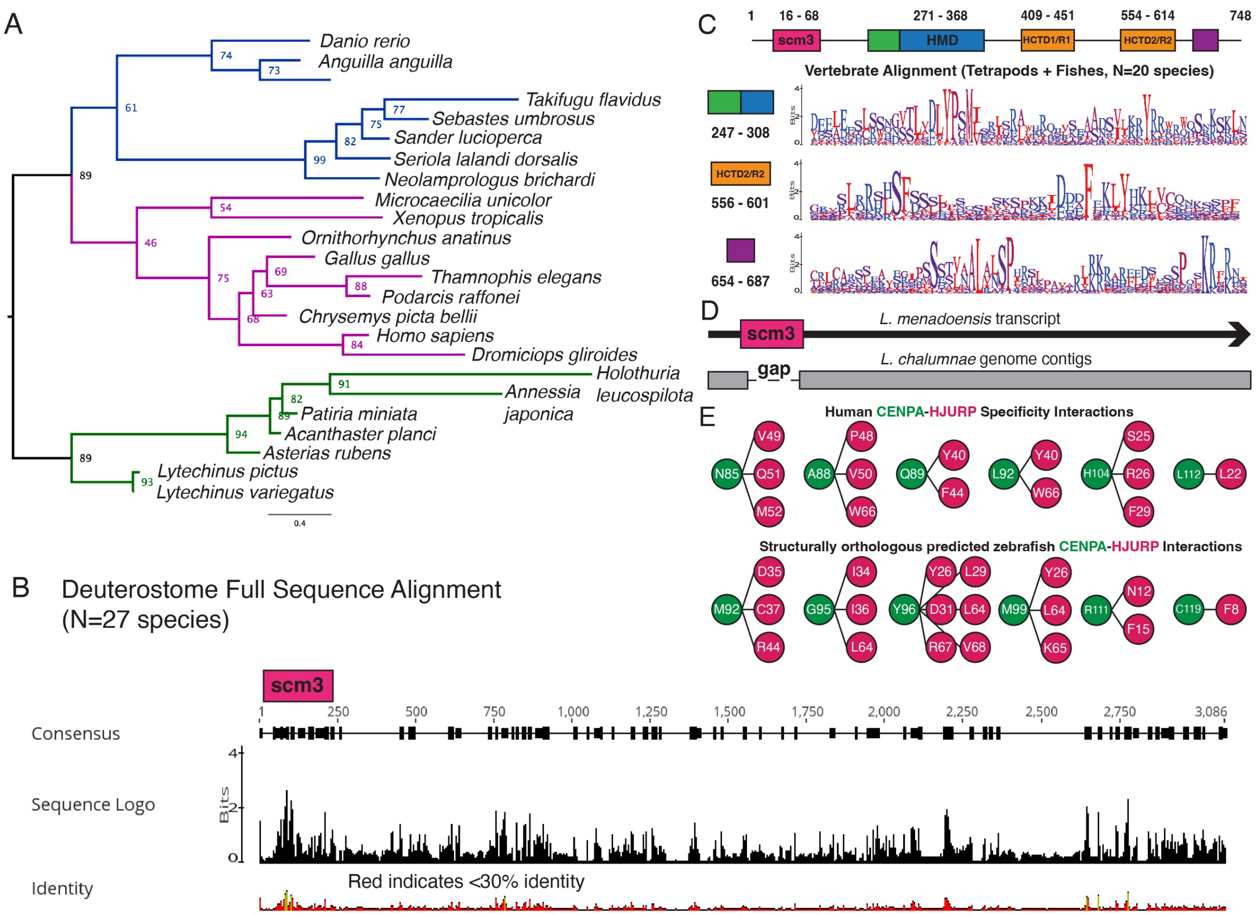


**Figure S4. Shared and diverged features of deuterostome HJURPs. A.** An expanded, duplicate phylogeny from Fig. 1 of representative deuterostome scm3 domains roughly follows the expected species tree, consistent with a monophyletic origin of HJURP in this clade. Nodes are presented with 1000 ultrafast bootstrap values. **C.** A sequence alignment of full-length putative deuterostome HJURP proteins shows generally poor conservation aside from the N-terminal scm3 domain region. **C.** A domain schematic for human HJURP with additional regions of shared homology between tetrapods and fish displayed as logo plots. Besides the previously identified HMD and HCTD2 domains, we find shared homology N-terminal to the HMD and close to the C terminus. **D.** The *L. chalumnae* genome likely contains a contig gap at the scm3 domain; alignments of translated contig regions to the *L. menadoensis* transcriptome shows coelacanths encode scm3 domain-containing HJURP orthologs. **E.** Structurally orthologous residues that contribute to CENPA-HJURP specificity differ almost entirely between zebrafish and human. Residues are shown that form physical interactions in either the human CENPA/H4/HJURP crystal structure or the predicted zebrafish CENPA/H4/HJURP structure.

A

| drHJURP Positively-correlated gene expression | |
|---|---|
| **Gene** | **r-value** |
| si:ch211-69g19.2 | 0.291 |
| tpx2 | 0.289 |
| cdk1 | 0.285 |
| plk1 | 0.283 |
| cenpf | 0.282 |
| cdc20 | 0.28 |
| ube2c | 0.28 |
| ccnb1 | 0.279 |
| mki67 | 0.279 |
| nusap1 | 0.278 |
| kpna2 | 0.277 |
| aspm | 0.273 |
| top2a | 0.273 |
| kifc1 | 0.271 |
| mad2l1 | 0.271 |
| kif11 | 0.268 |
| aurkb | 0.265 |
| aurka | 0.263 |

B

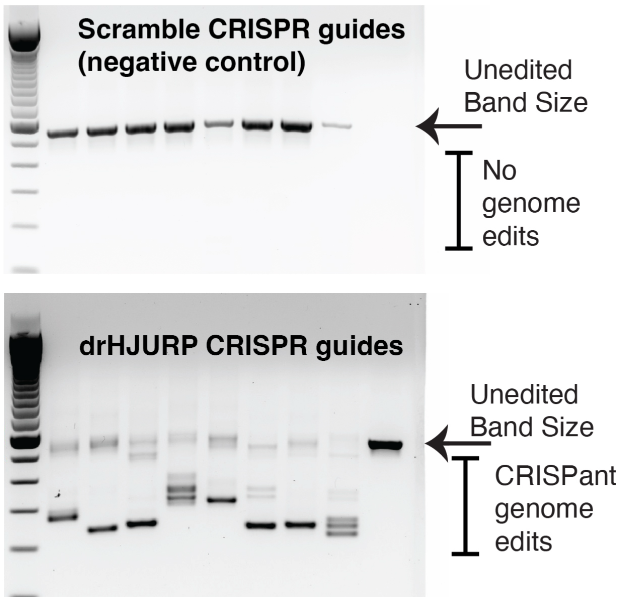


**Figure S5. Related to *in vivo* analysis of drHJURP. A.** The expression of the putative zebrafish HJURP is most positively correlated with primarily cell cycle genes such as cell cycle kinases (plk1, aurka/aurkb, cdk1) and spindle microtubule regulators (tpx2, aspm, kif11), among others. Data are provided via Daniocell[54]. **B.** Zebrafish CRISPR guides efficiently cause truncation at the HJURP locus (bottom) but non-targeting scrambled control guides fail to do so (top). Bands are PCR products amplified from the HJURP locus of individual CRISPant embryos; farthest right band is amplified from an uninjected control.

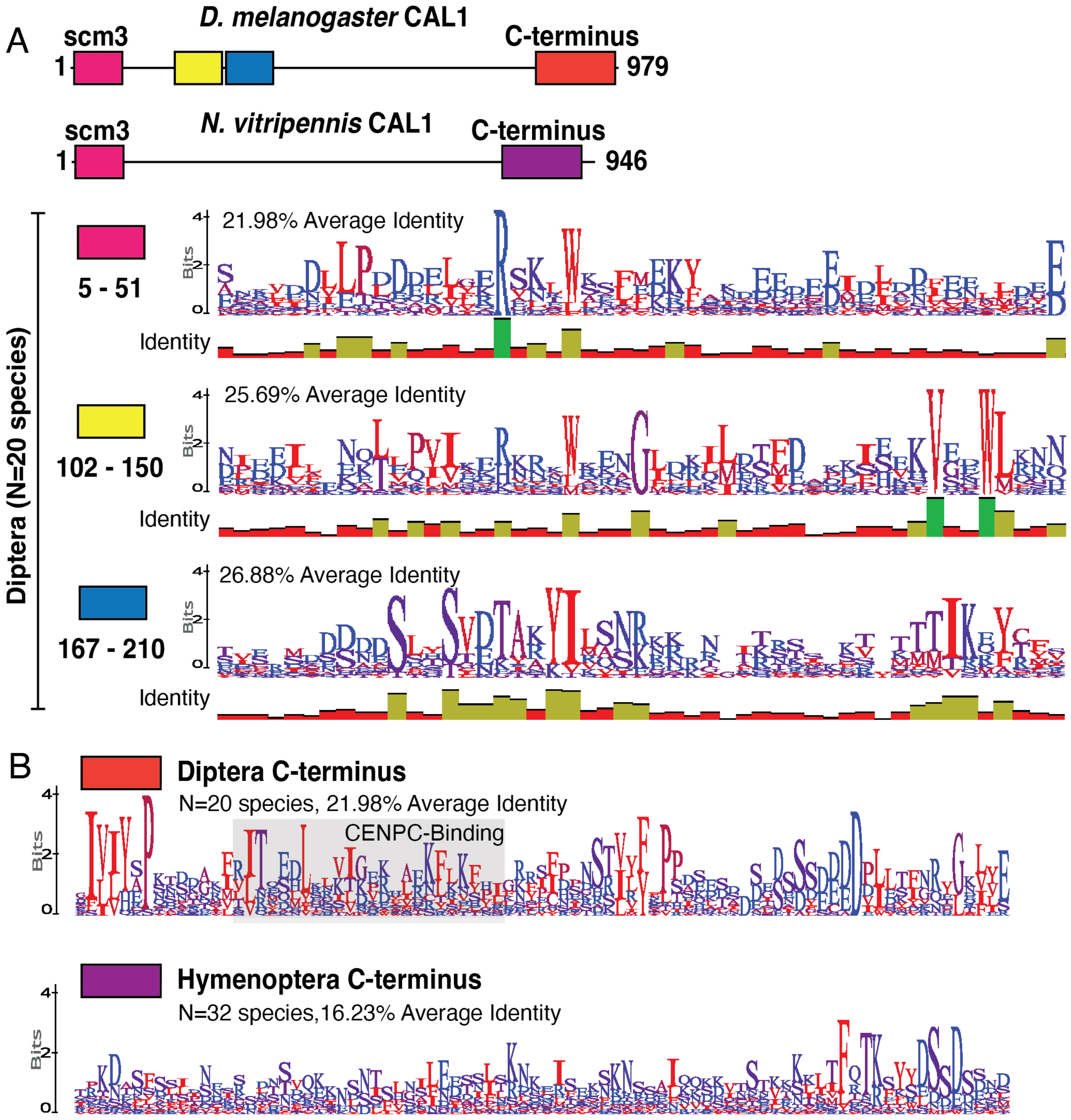


**Figure S6. Shared and diverged sequence features of CAL1. A.** A domain schematic for *Drosophila* and *Nasonia* CAL1 with additional regions of *Drosophila* homology beyond the scm3 domain displayed as logo plots. **B.** The Hymenopteran CAL1 C-terminus is less constrained than that of Dipteran CAL1.

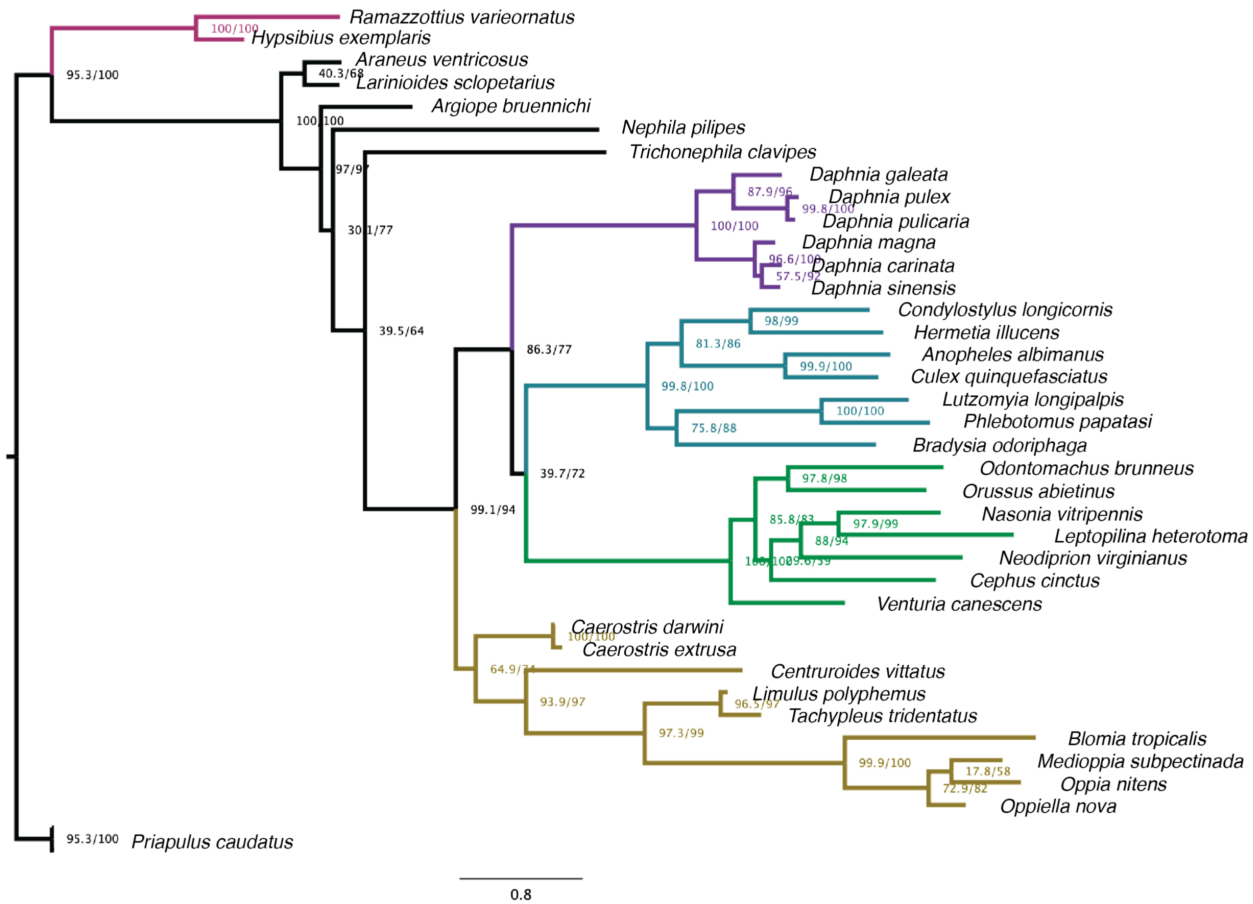


**Figure S7. An expanded phylogeny of panarthropod HJURP orthologs.** An expanded version of the tree in Fig. 3E generally follows the accepted species tree, although low homology and sparse sequence availability likely contributes to phylogenetic artefacts. Nodes are presented with 1000 ultrafast bootstrap/SH-aLRT replicate values.

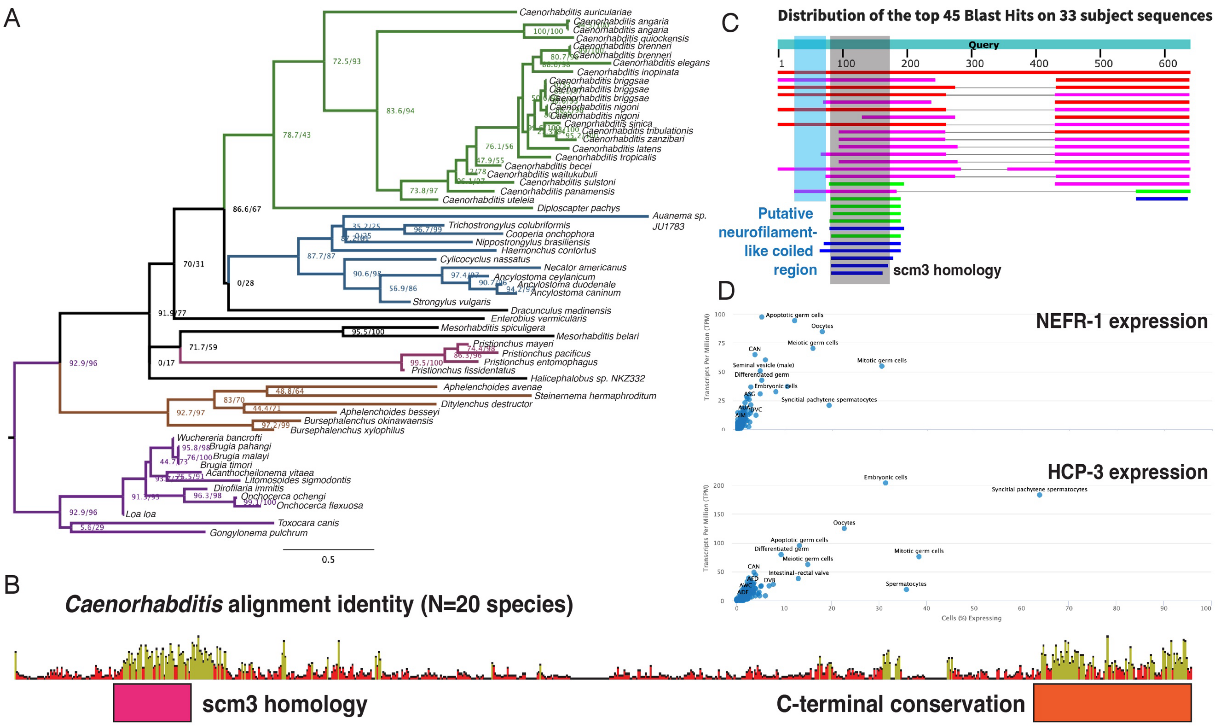


E

| GO term enrichment from positively-correlated gene expression (top 100 genes) | | |
|---|---|---|
| **GO term** | **p-value** | **Representative associated genes** |
| Cell cycle process | 1.28E-31 | KCA-1, SGO-1, PLK-1, FZY-1, CDC-25.1, POLE-1, KLP-19, MCM-6, ZIM-1, PLK-3 |
| Nucleus | 3.89E-31 | POLE-1, GON-4, MRE-11, ALG-5, BRD-1, EXO-1, NASP-2, GLP-1, ICP-1, CTF-4 |
| Nuclear division | 1.73E-30 | POLE-1, KCA-1, RDE-3, CDT-2, ZYG-1, KLC-1, GLD-3, ORC-1, CYB-2.2, FZY-1 |
| Sexual reproduction | 2.43E-30 | KCA-1, EGG-2, MARC-3, SPE-39, GLD-3, NOS-1, KLC-1, ZIM-1, RDE-3, PLK-1 |
| Intracellular organelle | 5.23E-30 | POLE-1, GON-4, MBTR-1, PASH-1, GNA-2, ZTF-1, RIF-1.1, VRK-1, ZTF-8, HIRA-1 |
| Organelle fission | 9.50E-30 | POLE-1, NOP-1, CTF-4, POLH-1, RDE-3, MCM-6, CYB-2.2, NOP-1, WDR-5.3, ORC-1 |
| Reproductive process | 1.37E-29 | KCA-1, GON-4, GNA-2, MARC-3, VRK-1, UNC-59, GLP-1, NOS-1, INX-22, ZIM-1, KLP-19 |
| Organelle | 1.56E-29 | POLE-1, PASH-1, TRUL-1, BRD-1, MUT-7, CITK-2, TAF-8, APTF-3, NPP-22, POLD-1 |
| Cell cycle | 2.65E-29 | KCA-1, IMA-2, CDT-2, ZYG-1, FZY-1, NOP-1, GLD-3, SZY-4, ICP-1, PLK-1, CDC-25.1 |

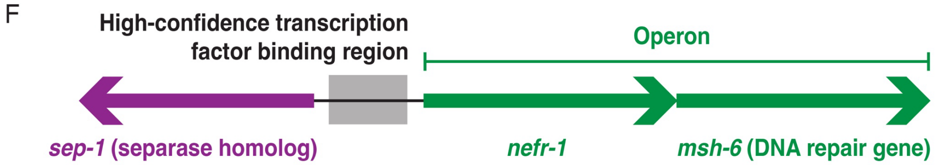


**Figure S8. NEFR-1 is a nematode cell cycle protein. A.** An expanded nematode phylogeny presented in Fig. 5B roughly follows the species tree, consistent with a monophyletic origin of HJURP$^{NEFR-1}$ in this clade. Nodes are presented with 1000 ultrafast bootstrap/SH-aLRT replicate values. **B.** Conservation pattern of NEFR-1 across *Caenorhabditis* shows there is relatively little sequence conservation outside of the N- and C-terminal regions. **C.** BLAST results show that the scm3 homology and C-terminal regions, rather than the putative neurofilament-like coiled coil region, are the most conserved within nematodes. **D.** NEFR-1, like the worm CENPA homolog HCP-3, is most highly expressed in germline dividing tissue such as oocytes, syncytial spermatocytes, and embryonic cells. Data are from single-cell expression profiles[68]. **E.** NEFR-1 has positive expression correlations with predominantly cell cycle genes. Data on the top 100 genes with positive expression correlation to *nefr-1* and their gene ontology (GO) analyses were provided via the WormBase[101] implementation of SPELL[124]. **F.** Schematic of the *nefr-1* syntenic neighborhood suggests *nefr-1* is expressed with other chromatin-associated cell cycle genes like *sep-1*, required for anaphase chromosome segregation, and *msh-6*, a DNA mismatch repair gene.

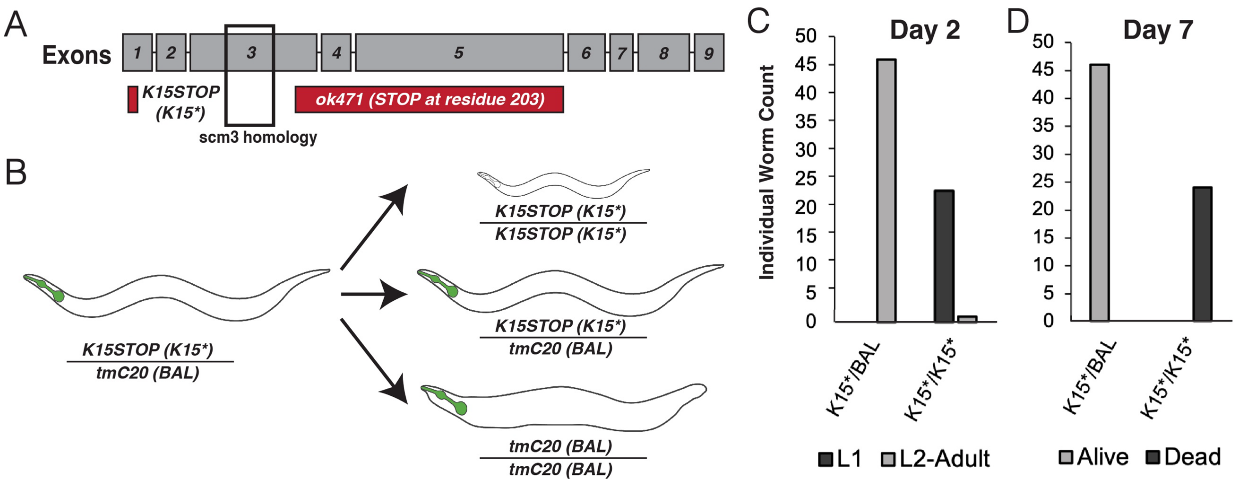


**Figure S9. NEFR-1 is an essential gene. A.** Gene structure of *nefr-1* shows that the previously characterized truncation allele *ok471* retains the scm3 domain, while the *K15** allele generated in this study, which includes a subsequent frameshift, produces a complete knockout. **B.** Balancer crossing scheme to assess viability of *nefr-1(K15*)* mutants. The *tmC20* balancer confers a GFP pharynx phenotype to heterozygotes and a GFP pharynx + dumpy phenotype to balancer homozygotes. **C.** Worms homozygous for *K15** fail to progress past the L1 stage and **D.** die within a week, while heterozygotes mature and survive into adulthood.

A

| GO term enrichment from top 500 mass spectrometry proteins (3xFLAG::NEFR-1) | | | |
|---|---|---|---|
| GO term | Fold Enrichment | p-value | Representative associated genes |
| Protein localization to kinetochore | 18.48 | 3.96E-06 | npp-10, czw-1, bub-3, rod-1, bub-1 |
| Protein-containing complex localization | 9.86 | 6.07E-04 | npp-10, lmn-1, bub-3, bub-1 |
| Mitotic cell cycle process | 8.06 | 2.10E-28 | spd-5, him-1, smc-3, mcm-4, tba-2, zyg-9, plk-1, hcp-2 |
| Mitotic DNA replication | 24.64 | 6.66E-07 | mcm-4, mcm-6, mcm-2, mcm-1, pola-1 |
| Nuclear cell cycle DNA replication | 44.35 | 2.55E-07 | mcm-4, mcm-2, mcm-3, pola-1 |
| Regulation of protein localization to the ER | 29.57 | 1.50E-03 | icd-2, icd-1 |
| Negative regulation of protein localization | 7.39 | 1.89E-03 | icd-2, icd-1, ftt-2, cdc-48.2 |
| Meiotic cell cycle process | 6.95 | 6.69E-15 | klp-18, lin-5, cmd-1, plk-1, scc-3, top-2, hcp-1 |
| Anatomical structure morphogenesis | 2.55 | 6.44E-07 | myo-3, unc-70, let-418, unc-54, swsn-1, sma-1, hcp-2 |
| Tube development | 4.16 | 3.18E-04 | lam-1, hda-1, let-418, erm-1, pyr-1, spt-16 |
| Spindle assembly | 10.44 | 6.81E-07 | paa-1, klp-18, air-1, hcp-2, bmk-1, klp-7, hcp-1 |
| Meiotic spindle organization | 19.4 | 2.71E-08 | klp-18, tbb-2, zyg-9, hcp-1, hcp-2 |
| Meiotic chromosome segregation | 4.93 | 8.62E-05 | mix-1, plk-1, smc-4, dpy-28, bub-1 |
| Acyl-CoA metabolic process | 8.45 | 3.78E-06 | sucg-1, pdhb-1, dlat-1, acly-1, acly-2 |
| Tricarboxylic acid metabolic process | 31.05 | 3.20E-10 | aco-1, idhg-1, idh-1, cts-1, acly-1, acly-2 |

| GO term enrichment from top 500 mass spectrometry proteins (Negative control) | | | |
|---|---|---|---|
| GO term | Fold Enrichment | p-value | Representative associated genes |
| L-amino acid catabolic process | 13.85 | 1.27E-14 | dlst-1, got-2.1, ahcy-1, mccc-1, gdh-1 |
| Proteinogenic amino acid catabolic process | 12.85 | 4.86E-14 | dlst-1, got-2.1, got-2.2, F02A9.10, mccc-1 |
| Alanine catabolic process | 44.16 | 5.12E-04 | C32F10.8 |
| L-alanine metabolic process | 44.16 | 5.12E-04 | C32F10.8 |
| Regulation of protein localization to the ER | 44.16 | 5.12E-04 | icd-1, icd-2 |
| Negative regulation of protein localization | 11.04 | 1.24E-05 | icd-1, icd-2, cdc-48.2, ftt-2 |
| Ketone body metabolic process | 29.44 | 1.51E-03 | hphd-1, C05C10.3 |
| Fatty acid derived catabolic process | 29.44 | 1.51E-03 | hphd-1, C05C10.3 |
| Anatomical structure morphogenesis | 5.52 | 2.43E-07 | myo-3, unc-54, act-5, erm-1, unc-15, emb-8 |
| Tube development | 3.68 | 1.51E-03 | lam-1, hda-1, sma-1, let-418, pyr-1, spt-16 |
| Purine ribonucleoside triphosphate biosynthetic process | 14.13 | 5.02E-08 | asb-1, atp-1, T22D1.3, Y69A2AR.18, atp-2 |
| Purine ribonucleotide biosynthetic process | 7.18 | 2.75E-08 | gmps-1, atp-1, T22D1.3, Y69A2AR.18, adsl-1,adss-1 |
| GTP metabolic process | 29.44 | 1.51E-03 | T22D1.3, ndk-1 |
| AMP biosynthetic process | 26.49 | 1.12E-04 | adsl-1, adss-1, adk-1 |
| Mitotic spindle organization | 7.68 | 7.97E-06 | spd-5, paa-1, tpxl-1, let-92, cdc-48.2, dhc-1 |

B

| Enrichment of centromere-related genes | |
|---|---|
| Gene | Enrichment Ratio |
| nefr-1 | 121.41637 |
| hcp-3 | 13.26415418 |
| ska-3 | 26.52830837 |
| san-1 | 7.937375803 |
| ndc-80 | 6.581222193 |
| mis-12 | 6.452589214 |
| ska-1 | 5.732842823 |
| mdf-1 | 4.376921342 |
| rod-1 | 3.577183895 |
| knl-2 | 3.080405495 |
| fzy-1 | 2.955127307 |
| cls-2 | 2.845736138 |
| plk-1 | 2.81279933 |
| kbp-5 | 2.679961418 |
| klp-7 | 2.495102451 |
| hcp-6 | 2.349041227 |
| czw-1 | 2.335227449 |
| kbp-1 | 1.983611933 |
| smc-4 | 1.895932272 |
| hcp-4 | 1.727331268 |
| him-10 | 1.713225622 |
| bub-1 | 1.694628825 |
| mdf-2 | 1.686378382 |
| mel-28 | 1.641716674 |
| lis-1 | 1.504390021 |
| kbp-2 | 1.489298728 |
| mix-1 | 1.361745056 |
| knl-1 | 1.242814502 |
| hcp-1 | 1.17379193 |
| xwl-1 | 1.156735494 |
| hcp-2 | 1.136039254 |
| kbp-4 | 1.060791731 |
| bub-3 | 1.016521348 |
| knl-3 | 0.953266723 |
| kbp-3 | 0.880508413 |

C

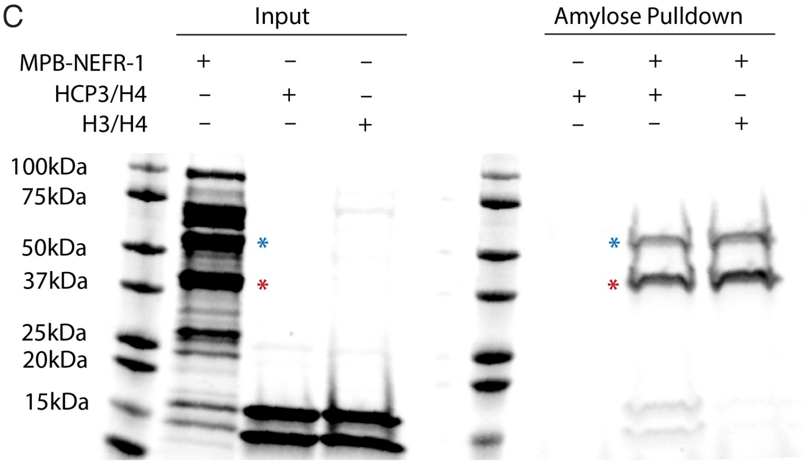


**Figure S10. NEFR-1 physically associates with CENPA$^{HCP-3}$. A.** Top 15 gene ontology (GO) analysis terms of mass spectrometry peptides identified in anti-FLAG immunoprecipitations from worm embryo lysate of either *3xFLAG::nefr-1* (left) or untagged control (right) strains. Tagged strains show specific enrichment of kinetochore and cell cycle-related genes. **B.** Normalized sum PEP score enrichment ratio of centromere-related genes[63] in *3xFLAG::nefr-1* immunoprecipitations over untagged controls. Genes highlighted in green, including CENPA$^{HCP-3}$ and NEFR-1, were identified exclusively in *3xFLAG::nefr-1* immunoprecipitations. **C.** Amylose pulldown assay with purified MBP-NEFR-1 and HCP-3/H4 or canonical H3/H4 tetramers. MBP-NEFR-1 interacts directly with HCP-3/H4 tetramers, as suggested by co-precipitation of species at HCP-3 and H4 sizes. Blue asterisk indicates MBP-NEFR-1; red asterisk indicates a non-NEFR-1 contaminant.

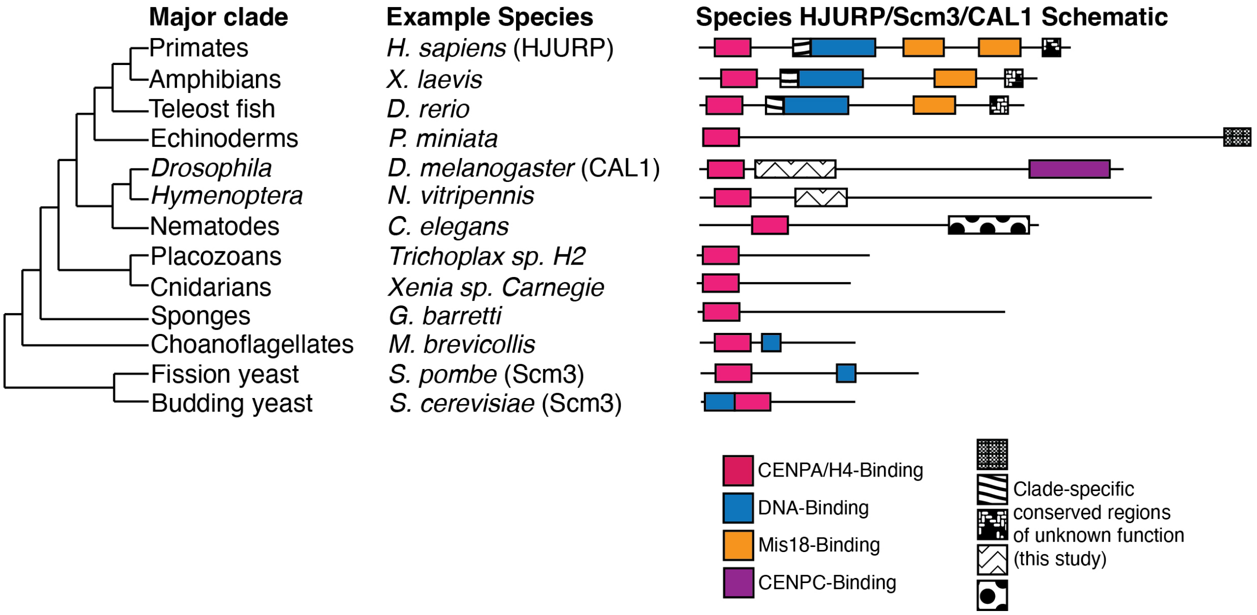


**Figure S11. scm3 domain-containing proteins are present in most metazoan clades.** We identify candidate HJURP orthologs across the metazoan tree and describe conserved domains previously unreported in these proteins that have yet to be functionally characterized.

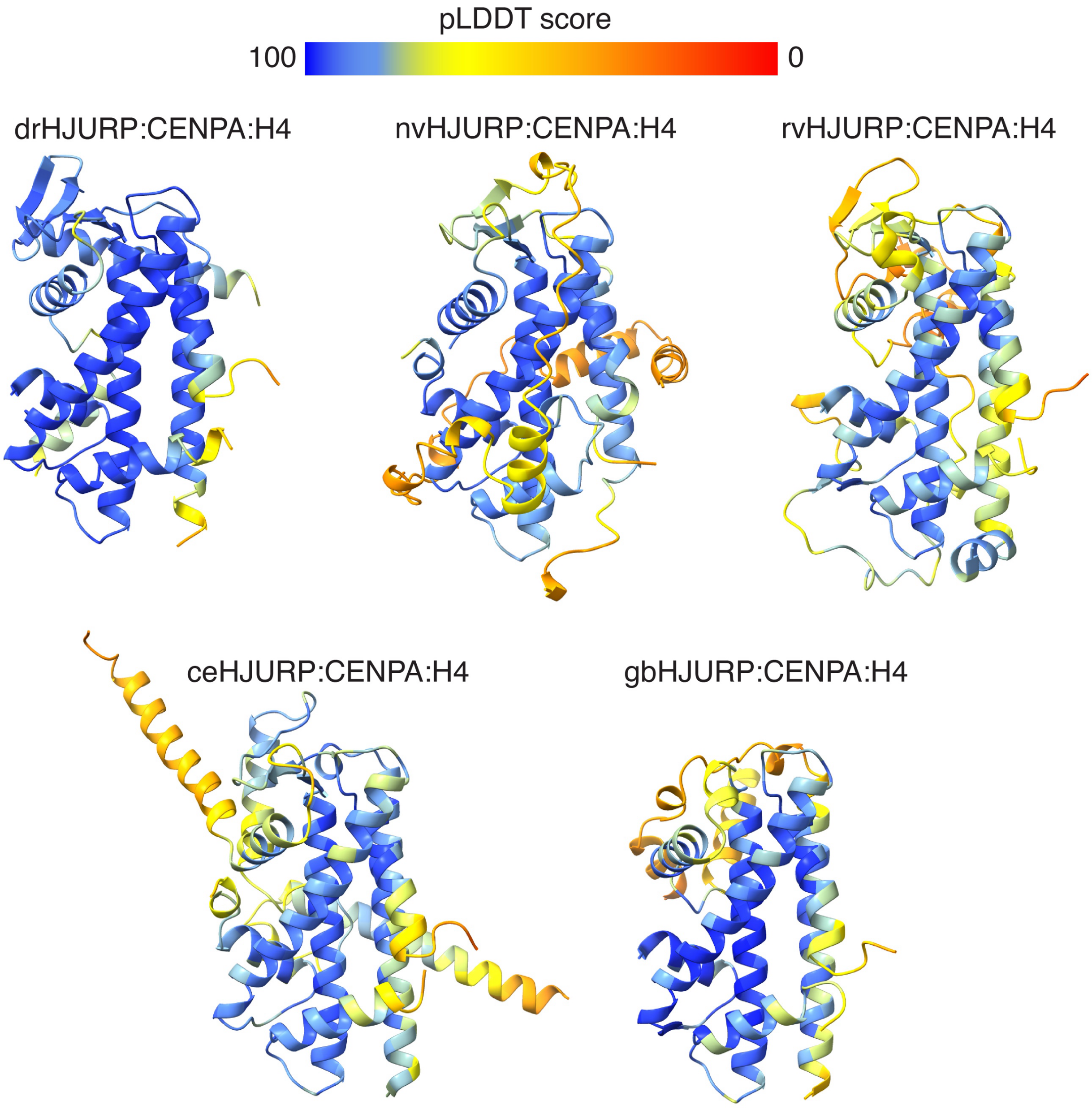


**Figure S12. AlphaFold predictions colored by pLDDT score.** All AlphaFold predictions presented in this study showed moderate to high confidence as measured by pLDDT (>80 at helical regions).

**Table S1.** A representative list of scm3 domain-containing proteins across the opisthokont lineage.

**Table S2.** Nucleic acid constructs used for this study.

**Table S3.** A list of *C. elegans* strains generated and/or used in this study.

**Table S4.** Sum PEP score ratios from 3xFLAG:NEFR-1 IP-MS data compared to the untagged control.

**Data S1.** Hidden Markov model (HMM) alignments used for this study.

**Data S2.** Protein alignments used for the generation of logo plots and trees presented in this study.

**Data S3.** Codon alignments used for the analyses in Figure 6E.

**Movie S1.** Timelapse of *D. rerio* embryo expressing GFP-CENPA and mCherry-HJURP shows discrete, cell cycle-dependent colocalization.
